# Modeling autism-associated SHANK3 deficiency using human cortico-striatal organoids generated from single neural rosettes

**DOI:** 10.1101/2021.01.25.428022

**Authors:** Yueqi Wang, Simone Chiola, Guang Yang, Chad Russell, Celeste J. Armstrong, Yuanyuan Wu, Jay Spampanato, Paisley Tarboton, Amelia N. Chang, David A. Harmin, Elena Vezzoli, Dario Besusso, Jun Cui, Elena Cattaneo, Jan Kubanek, Aleksandr Shcheglovitov

## Abstract

Our understanding of the human brain is limited by the lack of experimental models to mechanistically probe the properties of brain cells at different developmental stages under normal and pathological conditions. We developed a new method for generating human cortico-striatal organoids from stem cell-derived single neural rosettes (SNRs) and used it to investigate cortico-striatal development and deficits caused by the deficiency of an autism- and intellectual disability-associated gene *SHANK3*. We show that SNR-derived organoids consist of different cortico-striatal cells, including pallial and subpallial progenitors, primary cortical and striatal neurons, interneurons, as well as macroglial and mural cells. We also demonstrate that neurons in SNR-derived organoids are predictably organized, functionally mature, and capable of establishing functional neural networks. Interestingly, we found that the cellular and electrophysiological deficits in SHANK3-deficient SNR-derived organoids are dependent on the level of SHANK3 expression and that organoids with complete hemizygous *SHANK3* deletion have disrupted expression of several clustered protocadherins and multiple primate-specific zinc-finger genes. Together, this study describes a new method for using SNRs to generate organoids, provides new insights into the cell lineages associated with human cortico-striatal development, and identifies specific molecular pathways disrupted by hemizygous *SHANK3* deletion, which is the most common genetic abnormality detected in patients with 22q13 deletion syndrome.

## INTRODUCTION

Cortico-striatal networks have been implicated in numerous neurodevelopmental, psychiatric, and neurodegenerative disorders (Sanders et al., 2015; Shepherd, 2013; State and Šestan, 2012). However, our understanding of the cellular and molecular mechanisms underlying the development of these structures in the human brain remains limited. Most insights into cortico-striatal development have been obtained by studying the mouse brain; however, it is becoming increasingly clear that mouse and human brain development are different in terms of duration, produced anatomical structures, cellular composition, and underlying molecular pathways (Florio et al., 2017; Hodge et al., 2019; Lui et al., 2011; Onorati et al., 2014a; Rakic, 2009; Silbereis et al., 2016).

Studies on primary human brain tissue have provided new insights into the human-specific cytoarchitecture of the cerebral cortex and striatum (Fietz et al., 2010; Hansen et al., 2013, 2010; Ma et al., 2013; Onorati et al., 2014b; Paredes et al., 2016; Reillo et al., 2011) as well as the molecular programs associated with the development of these structures (Fan et al., 2018; Johnson et al., 2015; Kang et al., 2011; Krienen et al., 2019; Li et al., 2018; Miller et al., 2014; Nowakowski et al., 2017a; Onorati et al., 2014b; Pollen et al., 2015; Zhong et al., 2018). However, primary human brain tissue is not available from most patients and is not amenable to genetic manipulation for mechanistic studies.

To study the cellular and molecular mechanisms involved in human brain development and disorders, new methods have been pioneered for generating three-dimensional self-organized human neural tissue from pluripotent stem cells or multipotent neural stem cells – coined cerebral organoids or cortical spheroids (Eiraku et al., 2008; Kadoshima et al., 2013; Lancaster et al., 2013; Mariani et al., 2012; Paşca et al., 2015; Qian et al., 2016; Sloan et al., 2017; Watanabe et al., 2017). These studies have demonstrated that many important characteristics of human cortical development, including the specification of outer radial glia (oRG), formation of discrete cortical layers with inside-out organization, protracted neurogenesis, and consecutive gliogenesis can be recapitulated in stem cell-derived organoids. It was also shown that embryonic and induced pluripotent stem cell (PSC)-derived cerebral organoids have similar transcriptional and epigenetic profiles to primary fetal human neocortical tissue (Camp et al., 2015; Luo et al., 2016). Furthermore, different subtypes of subpallial medial ganglionic eminence (MGE)-derived inhibitory neurons (INs) and myelinating oligodendrocytes can also be generated in organoids, and different region-specific organoids can be fused together or transplanted to the mouse brain for studying interregional neuronal migration and anatomical and functional neuronal connectivity (Birey et al., 2017; Giandomenico et al., 2019; Madhavan et al., 2018; Mansour et al., 2018; Marton et al., 2019; Xiang et al., 2017, 2019). Importantly, organoids have also been produced from patient-derived and engineered stem cell lines and used to study the cellular and molecular deficits in human neurodevelopmental disorders associated with autism and malformations of cortical development, including microcephaly, lissencephaly, and tuberous sclerosis (Bershteyn et al., 2017; Birey et al., 2017; Blair et al., 2018; Lancaster et al., 2013; Mariani et al., 2015; Qian et al., 2016), highlighting the potential of brain organoids to revolutionize patient-specific drug discovery (Yang and Shcheglovitov, 2019).

Although organoids offer an enticing approach to modeling human brain development and diseases *in vitro*, the molecular pathways and cell lineages involved in the specification and differentiation of different human brain regions and cell types under normal and pathological conditions remain largely unknown. The major limitations hindering the use of organoids for basic research and in preclinical studies are their inconsistent cellular composition, unpredictable organization, and slow developmental maturation to study the functional properties of neurons and neural networks (Di Lullo and Kriegstein, 2017; Qian et al., 2016; Quadrato et al., 2017; Yang and Shcheglovitov, 2019).

Here, we describe a new method for generating organoids from stem cell-derived single neural rosettes (SNRs) (Elkabetz and Studer, 2008; Zhang et al., 2001; Ziv et al., 2015). We show that SNR-derived organoids consist of different cortico-striatal cells, including telencephalic pallial and subpallial neural progenitors (NPs), primary cortical excitatory and striatal inhibitory neurons, astrocytes, oligodendrocytes, and ependymal and mural cells. In addition, we demonstrate that SNR-derived organoids contain functionally mature neurons and neural networks capable of generating diverse electrical signals and oscillatory rhythms. Finally, we used SNR-derived organoids to investigate the cellular and molecular deficits caused by the deletions of the autism- and intellectual disability-associated gene *SHANK3* (Leblond et al., 2014; De Rubeis et al., 2014; Sanders et al., 2015). We show that SHANK3-deficient organoids have reduced numbers of excitatory synapses, elevated intrinsic excitability, and deficits in the expression levels of clustered protocadherins (Canzio and Maniatis, 2019; Lefebvre et al., 2012; Suo et al., 2012), such as *PCDHA6, PCDHGA10*, and *PCDHB15*, and primate-specific zinc finger (ZNF) genes, including *ZNF626, ZNF677, ZNF680, ZNF506*, and *ZNF229*. Interactive visualization of our transcriptomic and electrophysiological data is provided in our online browser (UBrain Browser: http://organoid.chpc.utah.edu). We propose that SNR-derived organoids can be used as a tractable platform for studying human cortico-striatal development and deficits associated with human neurodevelopmental disorders.

## RESULTS

### Generation of SNR-derived organoids

We used a modified dual-SMAD inhibition neural induction protocol (Chambers et al., 2009) and post-induction treatment with epidermal growth factor (EGF) and basic fibroblast growth factor (bFGF) to efficiently differentiate PSCs into neural rosettes (**Supplementary Fig. 1**). SNRs measuring 300–350 μm in diameter and composed of approximately 3000 cells with a clearly visible lumen were isolated for organoid generation (**Supplementary Fig. 2**). Cells in SNRs were characterized by expression of the telencephalic marker FOXG1, the NP markers PAX6 and SOX2, and proliferation markers KI67 and PH3 (**Supplementary Fig. 2d-j**). SNRs also contained few cells expressing the neuronal marker MAP2 (**Supplementary Fig. 2d-j**) and almost no cells expressing the subpallial MGE NP marker NKX2.1 (**Supplementary Fig. 2-3**). These results suggest that isolated PSC-derived SNRs primarily consist of proliferative NPs with dorsal telencephalic identities.

We generated SNR-derived organoids by propagating isolated SNRs in suspension culture for 2 weeks with EGF and bFGF to promote RG specification and proliferation (**Fig. 1a-c**). SNR-derived organoids demonstrated highly consistent growth across multiple batches, and a clearly visible internal lumen was observed in approximately 50% of organoids (**Supplementary Fig. 4**). Immunostaining with different cell-type specific antibodies (**Fig. 1d-i** and **Supplementary Fig. 5**) showed that most cells expressed the telencephalic marker FOXG1 (**Fig. 1d** and **Supplementary Fig. 5d**), with predominantly nuclear expression observed in NPs and both nuclear and cytoplasmic expression observed in neurons (Regad et al., 2007). The cells immediately adjacent to the lumen expressed NP markers SOX2 and PAX6, proliferation markers KI67 and PH3, and a RG marker phospho-vimentin (pVIM) (**Fig. 1d-g** and **Supplementary Fig. 5**), while the cells distributed more peripherally to the lumen expressed markers of both excitatory and inhibitory neurons, including MAP2, CTIP2, TBR1, and GAD67 (**Fig. 1e-g** and **Supplementary Fig. 5**). No cells expressed NKX2.1 (**Supplementary Fig. 5d**), and a relatively small proportion of cells expressed the marker of intermediate progenitors (IPs) TBR2 were found in 1-month-old SNR derived organoids (**Supplementary Fig. 6**). Mitotic RGs, unipolar cells co-expressing both PH3 and pVIM (∼0.5% of cells), were found on both the apical (∼60% of PH3- and pVIM-expressing cells) and basal sides (∼40% of PH3- and pVIM-expressing cells) of the area surrounding the lumen (**Fig. 1h**). Interestingly, both basal and apical RGs showed detectable expression of HOPX (**Supplementary Fig. 5)**, a specific marker of oRG at later neurodevelopmental stages (> GW16.7) and a non-specific RG marker at earlier stages (Nowakowski et al., 2016; Pollen et al., 2015; Thomsen et al., 2016), indicating an early developmental status of 1-month-old SNR-derived organoids. A similar non-specific HOPX expression pattern was observed in early (D56) stem cell-derived telencephalic organoids (Qian et al., 2016).

**Figure 1.**
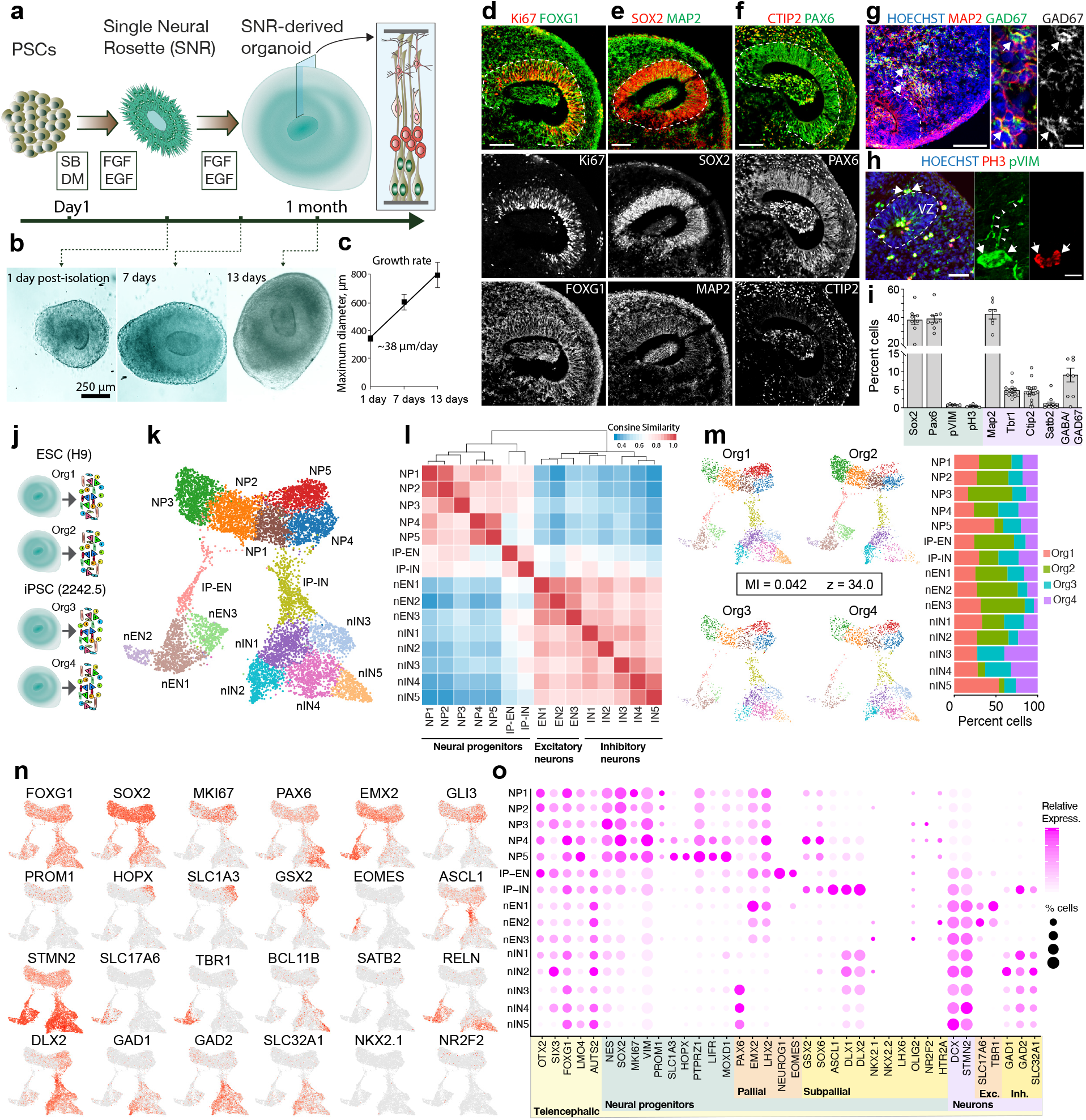
Reproducible generation of 1-month-old SNR-derived organoids containing multiple subtypes of telencephalic neural progenitors and excitatory and inhibitory neurons. **a**, Protocol schematic. **b**, Images of SNR-derived organoids at days 1, 7, and 13 post SNR isolation. **c**, Quantification of initial organoid growth (n = 32 rosettes obtained from 2 PSC lines). The line shows linear fit of the data. **d-h**, Images of organoid sections immunostained for FOXG1 and Ki67 (**d**), SOX2 and MAP2 (**e**), PAX6 and CTIP2 (**f**), MAP2 and GAD67 (**g**), and PH3 and pVIM (**h**). **i**, Quantification of cells expressing different cell type-specific markers (n = 6-19 organoids/1–10 sections per organoid). **j-k**, Single-cell RNA-seq experiment cartoon (**j**) and UMAP visualization of unsupervised cell clustering (n = 4 organoids) (**k**). **l**, Heatmap of relative gene expression similarity between cells in different clusters. **m**, Percentages of cells from different organoids in different cell clusters. **n**, Heatmap of expression profiles of different cell type-specific markers. **o**, Dot plot of expression profiles of different cell type-specific markers. Data are presented as mean ± standard error of the mean (s.e.m.). Scale bars = 50 and 10 μm (zoom-in). Abbreviations: NP, neural progenitor; IP, intermediate neural progenitor; EN, excitatory neuron; IN, inhibitory neuron.

### Reproducibility and characterization of cell types in 1-month-old SNR-derived organoids

To unbiasedly assess the reproducibility of our method for generating organoids, we performed single-cell RNA-sequencing (scRNA-seq) on 9577 cells collected from four 1-month-old SNR-derived organoids produced from two PSC lines in separate differentiation batches (**Fig. 1j**). Using unsupervised clustering, we identified 15 molecularly distinct cell clusters (**Fig. 1k-l** and **Supplementary Table 1**) and confirmed that each cluster was predominantly composed of cells collected from different organoids (**Fig. 1m**), suggesting a reproducible cellular composition was achieved with our approach. Indeed, the low mutual information (MI) between cluster assignment and sample identity, a statistical metric previously used for assessing organoid-to-organoid reproducibility based on scRNA-seq data (Velasco et al., 2019), supported this observation (MI = 0.042 and z-score = 34.0; lower scores indicate higher reproducibility). To broadly characterize the identities of cells in 1-month-old SNR-derived organoids, we investigated the expression of region- and cell-type specific markers in the identified clusters (**Fig. 1n-o, Supplementary Fig. 7**, and http://organoid.chpc.utah.edu). First, we found that cells in all clusters expressed the telencephalic marker *FOXG1* and showed enriched expression of dorsal anterior forebrain markers, including *OTX2, SIX3, LMO4*, and *AUTS2*. In addition, we observed no or undetectable expression of typical endodermal (*GATA4* and *GATA6*), mesodermal (*TBXT* [also known as BRACHYURY] and *HAND1*), or posterior and ventral brain markers, including *EN1, FOXA2, NKX2-1, NKX2-2, GBX2, DBX1, DBX2, HOXA2, HOXB2, HOXA1*, and *HOXB1* (**Supplementary Fig. 7** and http://organoid.chpc.utah.edu).

Second, we found that all cells could be divided into three groups: NPs (clusters: NP1, NP2, NP3, NP4, NP5, IP-EN and IP-IN), newborn excitatory neurons (nEN) (clusters: nEN1, nEN2, and nEN3), and newborn inhibitory neurons (nIN) (clusters: nIN1, nIN2, nIN3, nIN4, and nIN5) (**Fig. 1l**). Each group was characterized by the expression of well-known cell type-specific markers genes, including *NES, SOX2, MKI67, PAX6, EMX2*, and *LHX2* (NPs), *NEUROG1* and *EOMES* (IP-EN), *ASCL1, DLX1*, and *DLX2* (IP-IN), *SLC17A6* and *TBR1* (nENs), *and GAD1, GAD2, and SLC32A1* (or VGAT) (nINs) (**Fig. 1n-o**), and other genes with unknown or less-understood function in telencephalic development (**Supplementary Table 1** and http://organoid.chpc.utah.edu). The scRNAseq-based detection of both nENs and nINs in SNR-derived organoids is consistent with our immunostaining results (**Fig. 1i** and **Supplementary Fig. 5**) and confirms that both telencephalic ENs and INs are produced in SNR-derived organoids. Interestingly, cells in the NP5 cluster showed increased expression of typical oRG marker genes, including *SLC1A3*, HOPX, *PTRZ1, LIFR*, and *MOXD1* (Johnson et al., 2015; Pollen et al., 2015; Thomsen et al., 2016) (**Fig. 1o**), and cells in the NP4 cluster were characterized by enriched expression of typical lateral ganglionic eminence (LGE) markers, including *GSX2, SOX6, DLX1*, and *DLX2*, but not MGE or caudal ganglionic eminence (CGE) markers, such as *NKX2*.*1 and LHX6* or *NR2F2 and HTR2A*, respectively (**Fig. 1n-o**). These results suggest that 1-month-old SNR-derived organoids contain different subtypes of telencephalic NPs that contribute to the production of different subtypes of ENs and INs.

### Analysis of developmental lineages using RNA velocity

To gain insights into the specification of INs and different subtypes of NPs and neurons in 1-month-old SNR-derived organoids, we performed RNA velocity analysis using the velocyto algorithm (La Manno et al., 2018) (**Fig. 2a** and **Supplementary Fig. 8**). This algorithm predicts the future developmental states of individual cells using the relative abundance of spliced and unspliced transcripts, which can be used to identify the origin and destination states of the developmental progressions (**Supplementary Fig. 8a-c**). The RNA velocity analysis suggested that cells in the NP1 cluster are likely the origin cells that give rise to all other cells in 1-month-old SNR-derived organoids (**Supplementary Fig. 8b**). Interestingly, these cells show elevated expression of an apical RG marker *PROM1* (**Fig. 1n**) and several Wnt pathway-associated genes, including *RSPO1, GPC3*, and *FZD5* (**Fig. 2b**), suggesting that the Wnt signaling pathway may play an important role in regulating the properties of early telencephalic NPs in the human brain (Boyd et al., 2015).

**Figure 2.**
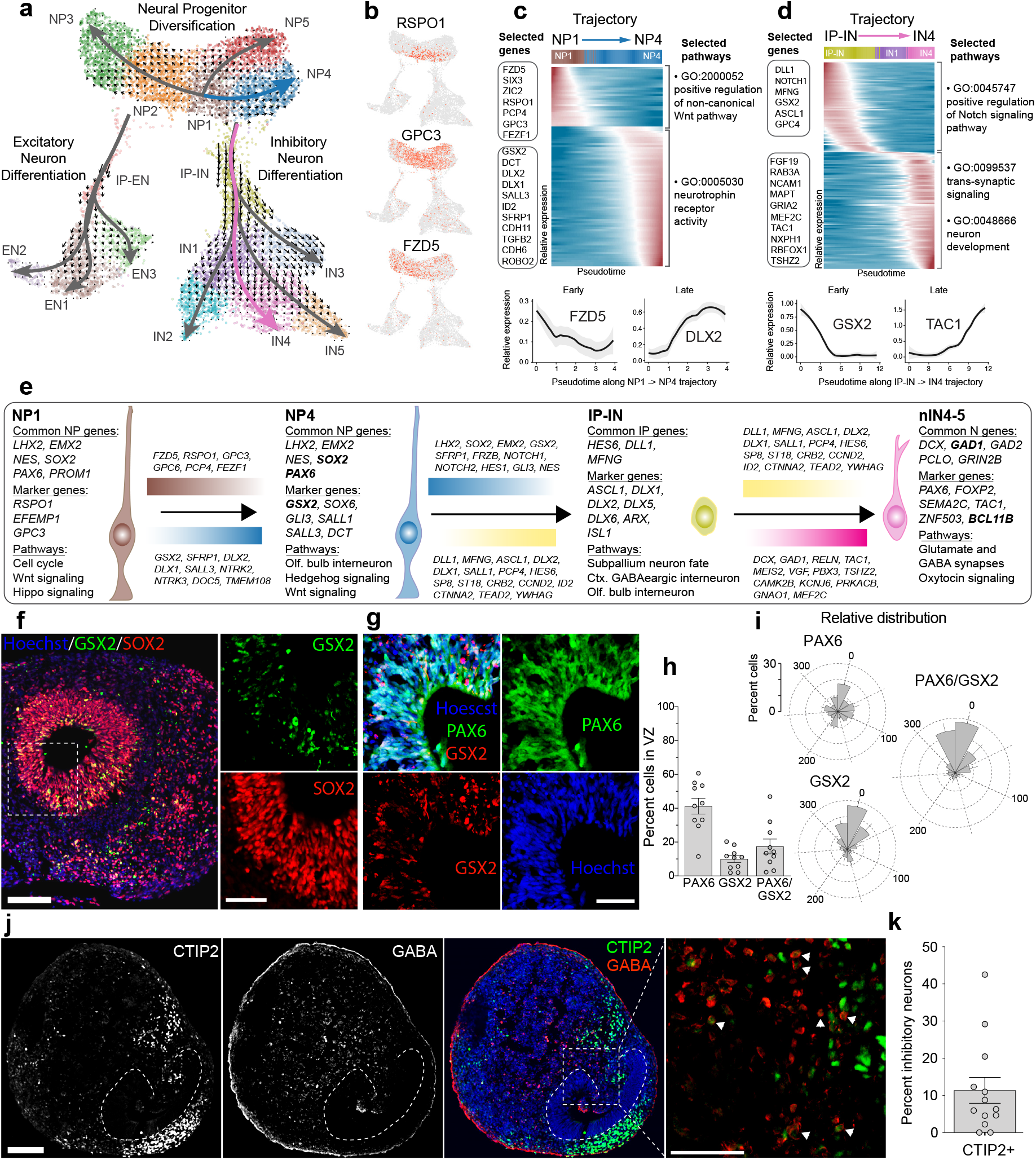
LGE origin of inhibitory neurons in 1-month-old SNR-derived organoids. **a**, RNA velocity vector fields and developmental trajectories. Small arrows indicate extrapolated future states of cells. **b**, Heatmap of the WNT signaling pathway genes enriched in NP1 (the origin cluster). **c-d**, Smoothed gene expression heatmaps of the top 150 genes differentially expressed along the NP1 -> NP4 (**c**) and IP-IN -> IN4 (**d**) trajectories. Genes are ordered by peak expression time on the pseudotime axis. Selected genes are listed on the left and shown at the bottom. The top gene ontology terms are listed on the right. **e**, Transcriptional programs and molecular pathways associated with specification of nIN4-5 in SNR-derived organoids inferred from the computational analysis. **f-g**, Organoid sections immunostained for GSX2 and SOX2 (**f**) and PAX6 and GSX2 (**g**). **h-i**, Percentages (**h**) and relative distributions (**i**) of PAX6-, GSX2- and PAX6/GSX2-expressing cells in the VZ-like area around the lumen (n = 10 organoids). **j**, Organoid sections immunostained for CTIP2 and GABA. **k**, Percentage of inhibitory neurons (GABA- or GAD67 expressing cells) co-expressing CTIP2. Data are means ± s.e.m. Scale bars = 100 (f, j) and 50 μm (zoom-in [**f**] and [**j**], g).

To infer the molecular programs associated with the specification of different lineages, we performed RNA velocity-guided lineage pseudotime inference analysis using Slingshot (Street et al., 2018) and identified genes that exhibit changing expression levels along different pseudotime trajectories (**Fig. 2a, Supplementary Fig. 8c** and **Supplementary Tables 2**). This analysis identifies the developmental trajectory of each cell type (**Fig. 2c-d, Supplementary Fig. 8d-g** and **Supplementary Fig. 9)**. For example, we found that IN lineage is likely specified through the transition along the NP1–>NP4 and IP-IN–>nIN trajectories (**Fig. 2c-d**). This transition is associated with expression changes for multiple genes, including those in the Wnt, Trk, Notch, Hippo, and Oxytocin signaling pathways (**Fig. 2e** and **Supplementary Tables 3**). As a result of the transition along the NP4–>IP-IN–>nIN4-5 trajectory, GSX2/PAX6/SOX2-expressing LGE-like NPs are converted into INs characterized by the co-expression of inhibitory and early striatal neuronal markers, including *GAD1, GAD2, FOXP2, TAC1, ZNF503* and *BCL11B* (also known as CTIP2) (Onorati et al., 2014a).

In the brain, striatal neurons are generated from a spatially regionalized pool of NPs in the dorsal LGE, which is adjacent to the pallial-subpallial boundary (PSB) and marked by unique co-expression of pallial marker PAX6 and subpallial marker GSX2 (Yun et al., 2001). To investigate the presence and distribution of LGE-like NPs and early striatal INs in 1-month-old SNR-derived organoids, we performed immunostainings for common LGE NP markers GSX2, SOX2, and PAX6 (**Fig. 2f-i**) and striatal neuron markers CTIP2, and GABA (**Fig. 2j-k**). We found that GSX2-expressing NPs were predominantly distributed to the VZ (∼15% of cells in the VZ) (**Fig. 2f-h**), while CTIP2-expressing INs (∼10% of all INs, **Fig. 2k**) were distributed in areas outside of the VZ (**Fig. 2j**). We found few to no DARPP32-expressing cells (not shown), suggesting an immature status of striatal INs in 1-month-old organoids. Interestingly, GSX2- and GSX2/PAX6-expressing cells showed preferential localization to a single area within the VZ, while PAX6-expressing cells were evenly distributed throughout the VZ (**Fig. 2i**). This suggests the presence of the pallial-subpallial regionalization in 1-month-old SNR derived organoids. A similar pattern of GSX2/PAX6-based regionalization was found in the pallial-subpallial region of fetal human brain tissue (PCW 11) (**Supplementary Fig. 10**). Together, these results confirm the results of RNA velocity analysis regarding the specification of nINs with early striatal identity in SNR-derived organoids and suggest that this analysis could be useful for inferring transcriptional programs associated with other developmental lineages in SNR-derived organoids.

We detected low to no expression of the superficial cortical layer marker *SATB2* (<1%), the mature striatal and cortical neuron maker *PPP1R1B* (also known as DARPP32), and the glial markers *GFAP, AQP4, SOX10*, and *MBP* (http://organoid.chpc.utah.edu), suggesting that 1-month-old SNR-derived organoids model a very early stage of human telencephalic development.

### Increased diversity of neural cell types in 5-month-old SNR-derived organoids

To encourage further development, we cultured SNR-derived organoids in Matrigel (MG) for an additional 4 months (**Supplementary Fig. 1**). MG-embedded organoids produced from different PSC lines and across multiple batches of differentiation consistently demonstrated growth and substantial expansion in size (**Fig. 3a-d**). To characterize the cellular composition of 5-month-old SNR-derived organoids and organoid-to-organoid reproducibility, we performed scRNA-seq and unsupervised clustering on 22,486 single cells collected from six SNR-derived organoids produced in separate batches from four different PSC lines (**Fig. 3e-h** and **Supplementary Table 1**). Notably, the sequenced organoids were composed of very similar cell types (**Fig. 3e-f**), with MI = 0.104 and z-score = 68.76. These scores were comparable to those observed for human and mouse cortical tissue samples (MI range: 0.008–0.064; z-score range: 2.2–41.4) and dorsally patterned 6-month-old organoids (MI = 0.089, z-score = 75.7)(Velasco et al., 2019), suggesting a high organoid-to-organoid reproducibility. Similar results for reproducibility were obtained using bulk RNA sequencing and correlation analysis (**Supplementary Fig. 11**), which is less sensitive for assessing cellular composition but frequently used in the previous studies on organoids (Luo et al., 2016; Qian et al., 2016; Watanabe et al., 2017; Yoon et al., 2019) and human fetal brain tissue (Ataman et al., 2016).

**Figure 3.**
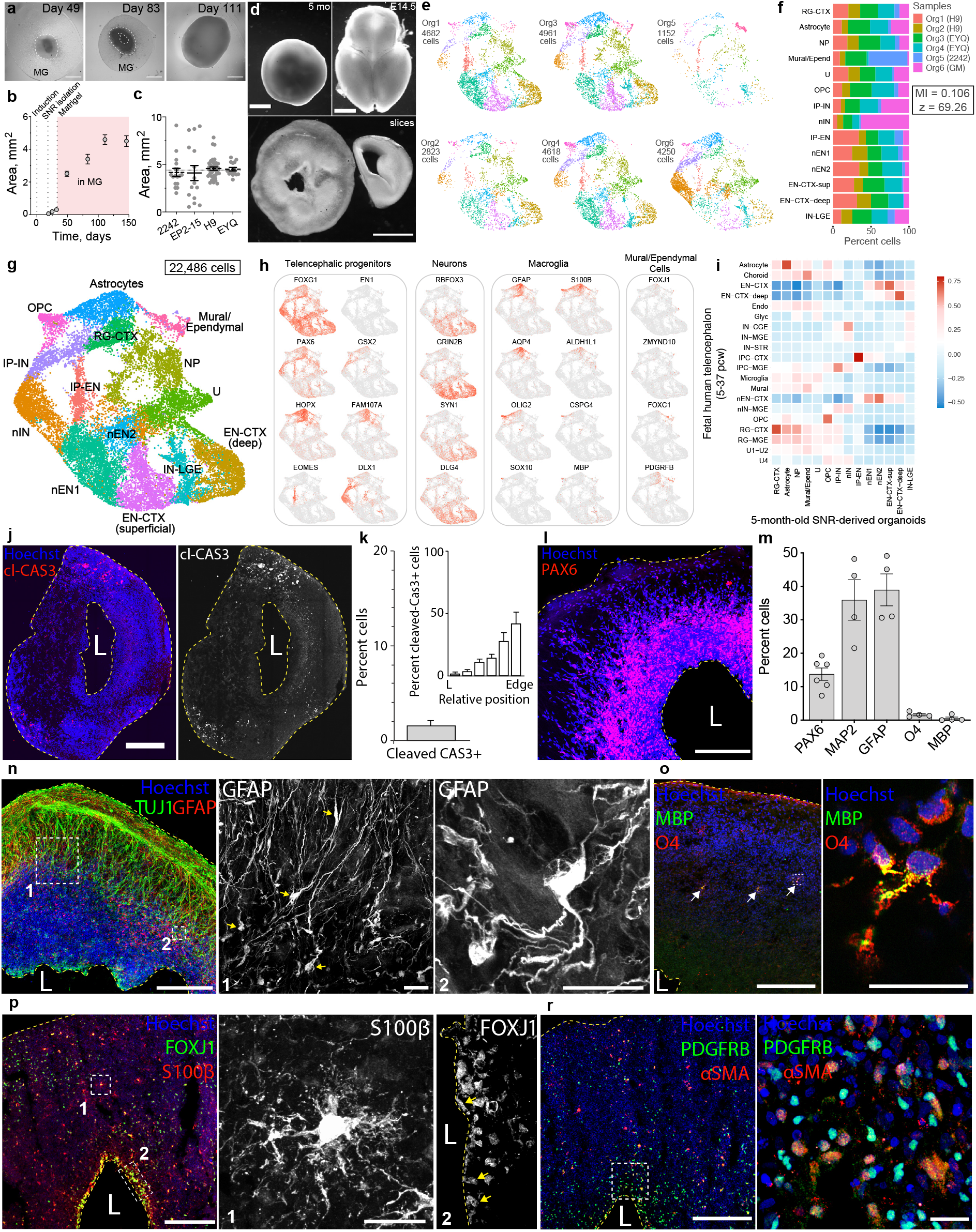
Diversity and organization of cell types in 5-month-old SNR-derived organoids. **a**, Images of SNR-derived organoids at day 49 (**a1**), 83, and 111 post SNR isolation. **b**, Quantification of organoid sizes over time (n = 25–47 H9 organoids from 2 differentiations). **c**, Quantification of sizes of 5-month-old SNR-derived organoids produced from different stem cell lines (n = 17–38 organoids). **d**, Images of a 5-month-old SNR-derived organoid (top) and 350-μm-thick organoid slice (bottom) together with E14.5 mouse brain (top) and coronal hemisphere slice (bottom) for relative visual size comparison. **e**, UMAP visualizations of unsupervised clustering of cells obtained from 5-month-old SNR-derived organoids generated from different stem cell lines. The clustering was performed on the combined dataset. **f**, Percentages of cells from different organoids in different clusters. **g**, UMAP visualization of combined 22,486 cells from 6 organoids. **h**, Heatmaps of expression of selected cell type-specific genes. **i**, Heatmap of Pearson correlation between cell clusters identified in 5-month-old organoids and those in primary human fetal tissue (Nowakowski et al., 2017). **j**, Organoid section immunostained for cleaved caspase-3. **k**, Percentage and distribution of cleaved caspase 3-expressing cells in organoid sections (n = 4 sections from 2 organoids). **l, n-r**, Organoid sections immunostained for PAX6 (**l**), TUJ1 and GFAP (**n**), O4 and MBP (**o**), FOXJ1 and S100β (**p**), and PDGFR-β and α-SMA (**r**). **m**, Percentages of cells expressing cell type-specific markers (n = 4–7 organoids/1–3 sections per organoid). Data are mean ± s.e.m. Scale bars = 1 mm (**a** and **d**), 500 (**j**), 200 and 20 (zoom-in) μm (**l** - **r**). Abbreviations: RG, radial glia; IP, intermediate neural progenitor; EN, excitatory neuron; IN, inhibitory neuron; CTX, cortex; OPC, oligodendrocyte progenitor cell; U, unknown.

To characterize the cellular composition of 5-month-old SNR-derived organoids, we investigated the expression of cell types-specific markers in different cell clusters (**Fig. 3g-h**). As in 1-month-old organoids, most cells in 5-month-old organoids showed robust expression of the telencephalic marker *FOXG1* (**Fig. 3h**) and no detectable expression of the endodermal, mesodermal, or posterior ectodermal markers (http://organoid.chpc.utah.edu), suggesting the telencephalic identity. In contrast to 1-month-old organoids, 5-month-old organoids contained cell clusters with increased expression of oRG markers *HOPX, FAM107A, TNC, PTPRZ1, LIFR*, and *MOXD1* (RG-CTX cluster, **Fig. 3h** and http://organoid.chpc.utah.edu); mature neuronal markers *RBFOX3* (NeuN), *GRIN2B, SYN1*, and *DLG4* (PSD95) (**Fig. 3h**); and macroglial markers *GFAP, S100B, AQP4, ALDH1L1, OLIG2, CSPG4* (NG2), *SOX10*, and *MBP* (**Fig. 3h**). Interestingly, we also found cells expressing the ependymal cell markers *FOXJ1, ZMYND10*, and *S100B* (Thomas et al., 2010; Zariwala et al., 2013) and the mural cell markers *FOXC1, PDGFRB*, and *ACTA2* (Vanlandewijck et al., 2018) (**Fig. 3h** and http://organoid.chpc.utah.edu). The gene expression profiles of different cell types in 5-month-old SNR-derived organoids were largely similar to those detected in cells from the fetal human cortex (PCW 5–37) (Nowakowski et al., 2017b) (**Fig. 3i)**, and a correlation analysis against human brain tissue at different developmental stages showed the strongest similarity to neocortical fetal brain at 12–24 PCW (**Supplementary Fig. 12**).

To characterize the distribution of the detected cell types, we performed immunostaining with different cell-type specific markers (**Fig. 3j-r)**. We confirmed that most cells in freshly dissociated organoids were viable (Trypan blue negative: 90±1.7%, n = 7 organoids) and that there was no enrichment of apoptotic, cleaved Cas3-expressing, cells near the lumen (**Fig. 3j-k)**. PAX6-expressing NPs (∼10% of all cells) were preferentially distributed around the lumen (**Fig. 3l-m**), while Tuj1- or Map2-expressing neurons (∼40%) and GFAP-expressing glia (∼40%) were positioned more basally to the lumen (**Fig. 3m-n)**. Interestingly, most GFAP-expressing cells were positioned away from the lumen and exhibited both oRG- and astrocyte-like morphologies (**Fig. 3n)**. Other cells, including S100®-expressing astrocytes (**Fig. 3p)**, O4- and MBP-expressing oligodendrocytes (**Fig. 3o)**, and PDGFR®- and ⟨ -SMA-expressing pericytes (**Fig. 3r**) were distributed throughout the organoid sections. Interestingly, the lumen in 5-month-old SNR-derived organoids was lined by S100®- and FOXJ1-expressing cells (**Fig. 3p**), suggesting their ependymal-like identity (Jacquet et al., 2009) and the idea that the lumen is a ventricle-like structure in SNR-derived organoids.

### Cortical and striatal neurons in 5-month-old SNR-derived organoids

To characterize the diversity of neurons in 5-month-old SNR-derived organoids, we analyzed the expression of different neuron-specific markers (**Fig. 4**). This analysis identified both ENs and INs in 5-month-old organoids (**Fig. 4a**, and http://organoid.chpc.utah.edu). ENs demonstrated increased expression of deep layer cortical markers *TLE4, LMO3, BCL11B*, and *FEZF2*, and superficial layer cortical markers *SATB2, CUX2, MDGA1*, and *FRMD4B*, suggesting their cortical identities. INs showed increased expression of IN markers *GAD1* and *GAD2* (GAD67 and GAD65, respectively), multiple striatal neuron markers, including *PBX3, ISL1, PPP1R1B (*DARPP*32), FOXP1, FOXP2, TAC1* (Substance P), *DRD2*, and *PENK* (Enkephalin) (Straccia et al., 2015); and the telencephalic interneuron markers, such as *SP8, NR2F2 (*COUP-TFII), *CALB2* (Calretinin), *VIP, CCK, HTR3A, SST, PVALB, CALB1, NPY, VIP, NOS1*, and *RELN* (**Fig. 4a** and http://organoid.chpc.utah.edu). Interestingly, a small proportion of cells in the IP-IN, IN, and IN-LGE clusters expressed NKX2.1 (http://organoid.chpc.utah.edu), which is consistent with the striatal identity of these neurons (Onorati et. al. 2014). In addition, some cells in the EN-CTX-deep cluster showed increased expression of late activity-regulated genes, such *OSTN* (Ataman et al., 2016), suggesting an advanced maturation status of neurons in 5-month-old SNR-derived organoids.

**Figure 4.**
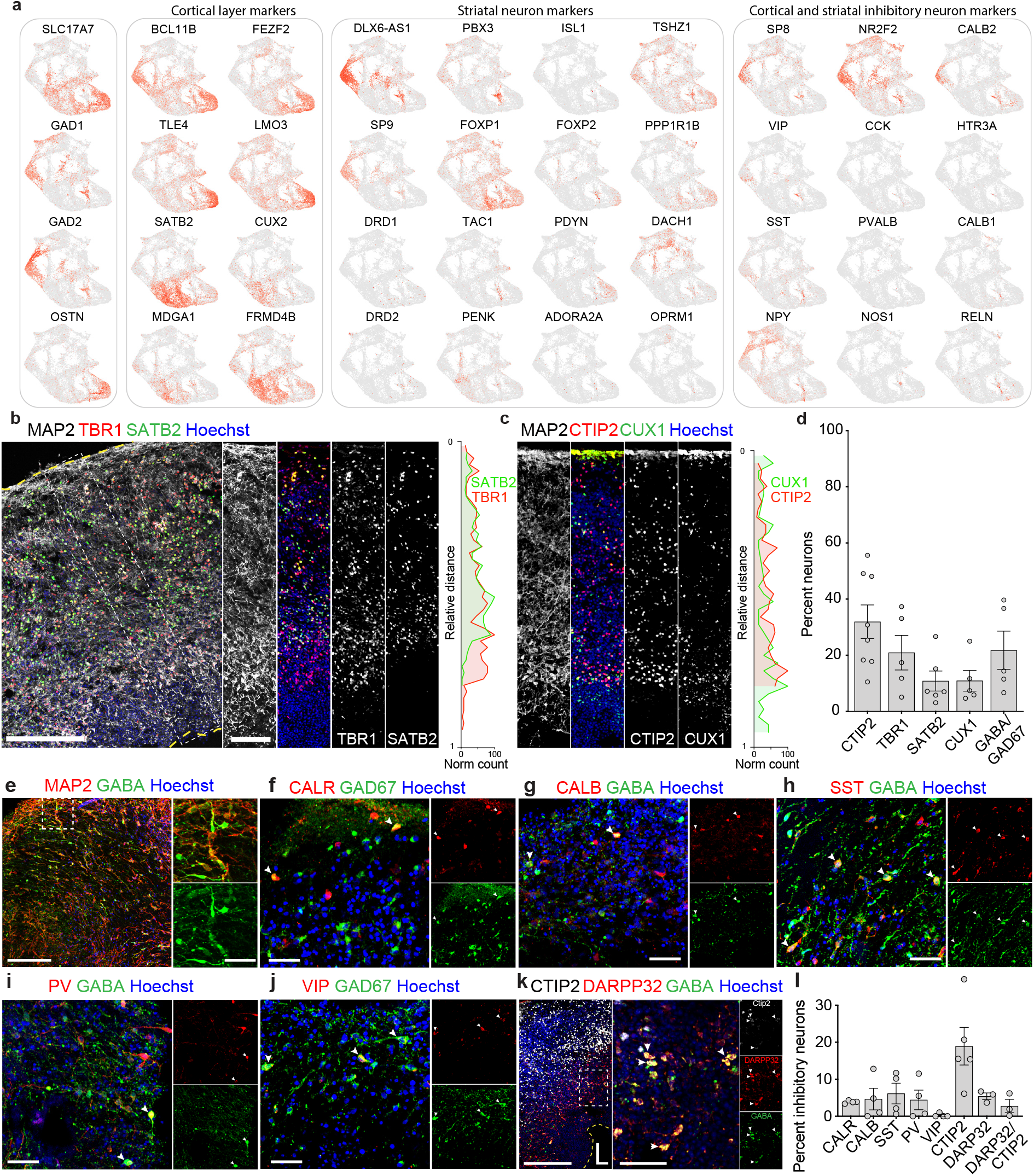
Neuronal diversity in 5-month-old SNR-derived organoids. **a**, Heatmap of expression of selected neuronal marker genes. **b-c**, Organoid sections immunostained for MAP2, TBR1, and SATB2 (**b**) and MAP2, CTIP2, and CUX1 (**c**) and relative distributions of TBR1- and SATB2-(**b**) and CTIP2- and CUX1-expressing cells (**c**). **d**, Quantified expression of cell type-specific markers in organoid sections (n = 6–8 organoids/1–2 sections per organoid). **e-k**, Organoid sections immunostained for MAP2 and GABA (**e**), calretinin and GAD67 (**f**), calbindin and GABA (**g**), somatostatin and GABA (**h**), parvalbumin and GABA (**i**), VIP and GAD67 (**j**), and CTIP2, DARPP32, and GABA (**k**). **l**, Quantification of inhibitory neuron subtypes (n = 3–5 organoids/1–3 sections per organoid). Data are presented as mean ± s.e.m. Scale bars = 200 and 50 (zoom-in) μm.

We next investigated neuron distribution within the organoids by immunostaining (**Fig. 4b-l**). Neurons co-expressing MAP2 and the deep layer cortical marker TBR1 or CTIP2 (20-40% of all neurons [**Fig. 4d**]) were predominantly distributed in the deeper layers in organoid sections (**Fig. 4b-c**), while neurons co-expressing MAP2 and the superficial layer cortical marker SATB2 (∼10% of all neurons [**Fig. 4d**]) resided more superficially (**Fig. 4b**). These distributions are consistent with a laminar organization of cortical neurons in organoid tissue. We found no layer-specific distribution for CUX1-expressing cells (**Fig. 4c-d**). In contrast to ENs, INs (∼20% of all neurons [**Fig. 4d-e**]) were distributed throughout the tissue (**Fig. 4e-k**). We could distinguish several subtypes of inhibitory interneurons, including those expressing GAD67 and calretinin (∼4%, **Fig. 4f**), GABA and calbindin (∼5%, **Fig. 4g**), GABA and somatostatin (∼7%, **Fig. 4h**), GAD67 and parvalbumin (∼4%, **Fig. 4i**), and GAD67 and vasoactive intestinal polypeptide (VIP) (∼4%, **Fig. 4j**). Interestingly, GABAergic neurons co-expressing the striatal markers CTIP2 (∼20%), DARPP32 (∼5%), or both CTIP2 and DARPP32 (∼3%) (**Fig. 4l**) were primarily found beneath the cortical layers (**Fig. 4k**), resembling the relative organization of the cortex and striatum in the telencephalon. These results suggest that SNR-derived organoids could be used for modeling human cortico-striatal development.

### Functional neurons and networks in SNR-derived organoids

To investigate the functional properties of neurons in 5-month-old SNR-derived organoids, we performed patch-clamp electrophysiology on acute slices. We targeted cells in the peripheral areas that showed layered organization and contained neurons with pyramidal-like morphology, complex dendrites, and dendritic spines (**Fig. 5b**). Remarkably, about 86% (103/120) of recorded cells fired action potentials (APs) and generated diverse patterns of electrical activity (**Fig. 5c** and **Supplementary Table 4)**. To unbiasedly characterize the diversity of functional neurons in 5-month-old SNR-derived organoids, we extracted multiple electrophysiological characteristics from the recordings of voltage deflections in response to different somatic current injections and performed unsupervised clustering (**Fig. 5d**). All recorded cells could be divided into five groups based on their passive membrane properties (input resistance, resting membrane potential, and capacitance), single action potential (AP) profile (AP amplitude, latency, threshold, width, after-hyperpolarization potential amplitude [AHP], and sag) and AP firing pattern (frequency and adaptation) (**Fig. 5d**). Group 2 cells showed the least mature functional properties, including high input resistance, depolarized resting membrane potential, and wide AP (**Fig. 5c2** and http://organoid.chpc.utah.edu), while cells in groups 1 and 3–5 were more mature (**Fig. 5c1,3-5** and http://organoid.chpc.utah.edu) and capable of firing multiple APs without adaptation (group 1), with a sag (group 3), with strong AP adaptation (group 4), or with relatively hyperpolarized resting membrane potentials and depolarized threshold for AP generation (group 5). These results indicate that 5-month-old SNR-derived organoids contain a diverse population of functional neurons.

**Figure 5.**
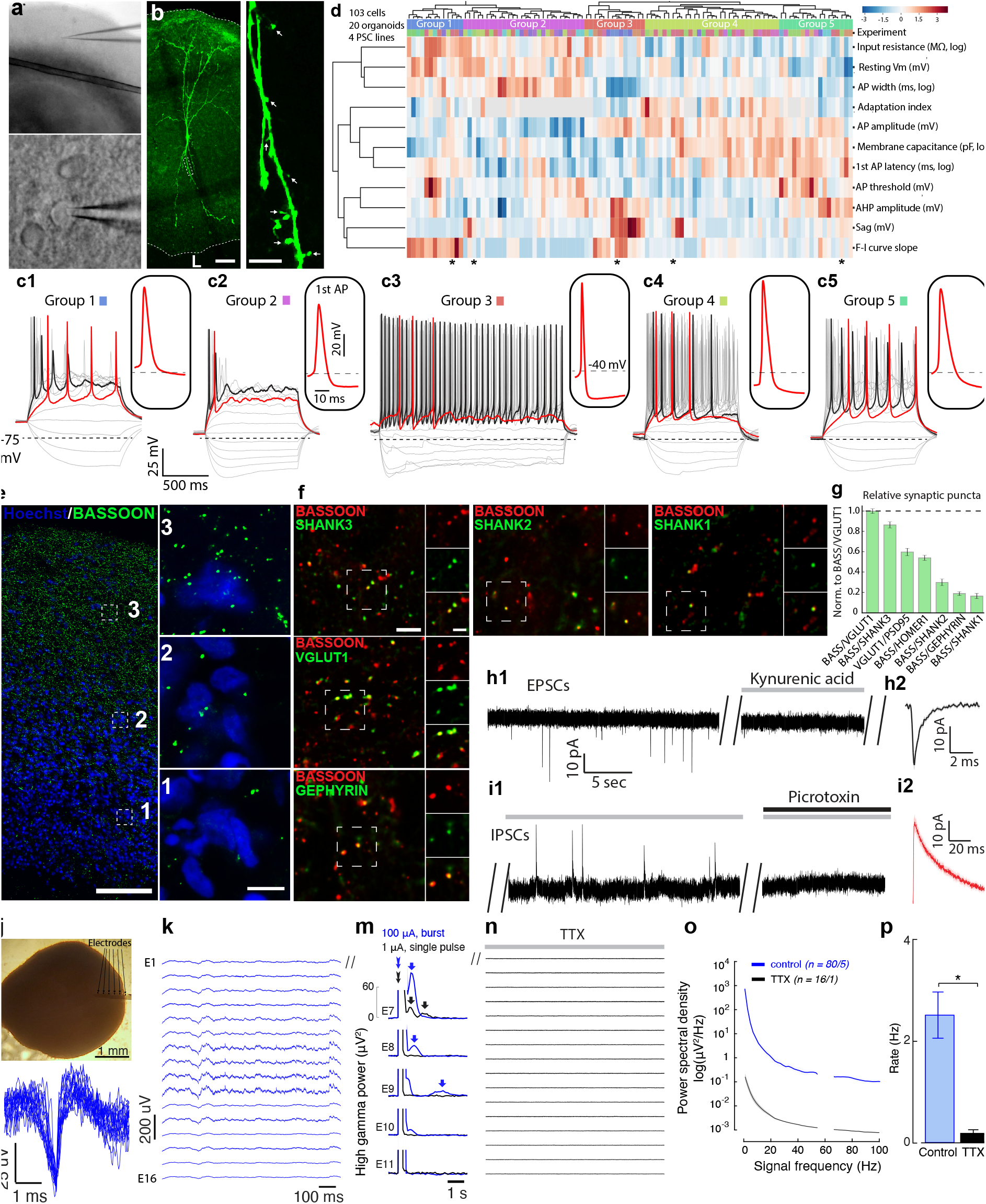
Functional neurons and neural networks in 5-month-old SNR-derived organoids. **a** Low-(top) and high-(bottom) resolution images of a 5-month-old organoid slice used for electrophysiology. **b**, Image of a biocytin-filled control neuron in organoid slice after electrophysiology. **c**, Traces of membrane potentials recorded from cells (marked with asterisk in **d**) in different clusters in response to Δ5 pA somatic current injections. **d**, Heatmap of unsupervised hierarchical clustering of recorded neurons based on non-redundant electrophysiological characteristics (n = 103 cells from 4 2242-5, 3 EP2-15, 11 H9, and 2 EYQ organoids). **e**-**f**, Organoid section immunostained for presynaptic marker BASSOON. (**e**), BASSOON and SHANK3, SHANK2, SHANK1, VGLUT1, or GEPHYRIN (**f**). **g**, Quantification of excitatory and inhibitory synaptic puncta (n = 8–28 sections from 1–3 control organoid sections). **h-i**, Traces of spontaneous excitatory postsynaptic currents (EPSCs) (**h1-2**) and inhibitory postsynaptic currents (IPSCs) (**i1-2**) recorded at different holding membrane potentials without inhibitors and with 3 mM kynurenic acid (glutamate receptor antagonist) or 100 μM picrotoxin (GABA receptor antagonist). Average EPSC (**i2**) and IPSC (**i2**) traces. **j**, Image of a 5-month-old SNR-derived organoid implanted with a 16-channel multi-electrode array. **k-n**, Spontaneous electrical activity recorded from different electrodes (E1–16) before (**k**) and after tetrodotoxin (TTX, 1 µM) application (**n**). Waveforms extracted after filtering of low-frequency components with a 300-Hz high-pass filter (**l**). Extracellular potentials recorded from different electrodes after delivery of stimuli of varying intensity at E7 (**m**). **o**-**p**, Spectral analysis of recorded signals (**o**) and average rate of high-frequency events (> 300 Hz) before (n = 80 fields/5 organoids) and after TTX application (n = 16 fields/1 organoid) (**r**). Data are mean ± s.e.m.; *P<0.05, Mann-Whitney test. Scale bars = 50 and 10 (zoom in) (**b**), 200 and 20 (zoom in) (**e**), and 20 and 10 (zoom in) (**f**) μm.

We next investigate the synaptic properties of neurons in 5-month-old SNR-derived organoids via synaptic immunostaining and physiology experiments (**Fig. 5e-i**). Interestingly, the density of synapses in organoid sections, visualized by immunostaining for presynaptic marker BASSOON (**Fig. 5e**), increased along the lumen to the periphery axis, suggesting increased maturation of cells at the periphery compared with those at the lumen. Co-immunostaining with different pre- and postsynaptic markers, including BASSOON/VGLUT1, BASSOON/SHANK3, VGLUT1/PSD95, BASSOON/HOMER1, BASSOON/SHANK2, BASSOON/SHANK1 and BASSOON/Gephyrin (**Fig. 5f-g**), confirmed the presence of different subtypes of putative excitatory and inhibitory synapses in 5-month-old SNR-derived organoids. Interestingly, SHANK3-containing synaptic puncta were more abundant than those containing SHANK2 or SHANK1 (**Fig. 5g**), suggesting preferential expression of SHANK3 at early excitatory synapses in human telencephalic tissue.

To investigate functional synapses, we performed patch-clamp electrophysiology on organoid slices (**Fig. 5h-i**). Both spontaneous excitatory and inhibitory postsynaptic currents (EPSCs and IPSCs, respectively) were detected in ∼56% of recorded cells (9 out of 16) (**Fig. 5h-i** and **Supplementary Table 4**). EPSCs were characterized by fast rise and decay kinetics (**Fig. 5h2**) and could be completely blocked by application of the AMPA and NMDA receptor antagonist, kynurenic acid (**Fig. 5h1**). IPSCs showed significantly slower kinetics (**Fig. 5i2**) and could be completely abolished by application of a GABA-A receptor antagonist, picrotoxin (**Fig. 5i1**). These results demonstrate that human neurons in 5-month-old SNR-derived organoids are synaptically mature and interconnected through both excitatory and inhibitory synapses.

To test whether organized and functional neurons in 5-month-old SNR-derived organoids establish functional neural networks, we implanted flexible miniature 16-channel multi-electrode arrays in 5-month-old SNR-derived organoids (**Fig. 5j**). Interestingly, 1-month post-implantation, we observed clear patterns of spontaneous electrophysiological activity with waveforms resembling typical extracellularly recorded APs (**Fig. 5k-l**). To identify functional neural connections, we electrically stimulated one active electrode and recorded the responses from all electrodes (**Fig. 5m**). We found that relatively strong stimuli (100-mA bursts) activated not only the stimulated channel but also elicited a sequence of responses in the adjacent electrodes. Weaker stimuli (1-mA single pulse) did not induce responses in the adjacent electrodes, which could be due to incomplete maturation of functional neural networks in 6-month-old SNR-derived organoids. Importantly, the observed activity was abolished by bath application of a sodium channel antagonist, tetrodotoxin (TTX) (**Fig. 5n**), suggesting the AP-dependent nature of the detected functional connections.

To determine whether oscillatory rhythms are generated in SNR-derived organoids, we performed a spectral analysis of the recorded signals, before and after TTX administration (**Fig. 5o**). In all recorded organoids, we observed a 1/f spectrum, reminiscent of a human electroencephalogram (EEG) (Trujillo et al., 2019). However, unlike a human EEG, we did not observe peaks in the alpha or beta bands, which are associated with the activity of thalamic neurons and thalamo-cortical networks. Oscillations detected in SNR-derived organoids appeared at a rate of ∼2.5 Hz, which substantially exceeded the noise level, and were abolished upon TTX application (**Fig. 5p**). Together, these results indicate the presence of functional rhythmically oscillating neural networks in SNR-derived organoids.

### Deficits in SHANK3-deficient SNR-derived organoids

To test whether SNR-derived organoids can be used to model neurodevelopmental disorders, we generated CRISPR/Cas9-engineered PSC lines with partial homozygous or complete hemizygous *SHANK3* deletions (**Fig. 6a-b**). The homozygous *SHANK3* deletion resulted in complete loss of expression of all SHANK3 isoforms (**Supplementary Fig. 13**), whereas complete hemizygous *SHANK3* deletion, which recapitulates the genetic abnormality detected in most Phelan-McDermid syndrome patients (Wilson et al., 2003), resulted in ∼50% loss of expression of the longest and shortest SHANK3 isoforms (Chiola et al., 2021). Interestingly, 1-month-old *SHANK3*- /- organoids were significantly smaller than isogenic control (iCtrl) organoids and contained a significantly reduced proportion of neurons that were also smaller than those in iCtrl) organoids (**Supplementary Fig. 14**). The size and neuron proportion phenotypes, however, were not detected in 1-month-old *SHANK3*+/- organoids (**Supplementary Fig. 14**), suggesting that the level of SHANK3 expression may influence the severity and/or manifestation of deficits. We observed no overt size deficits in 5-month-old SHANK3-deficient organoids (**Fig. 3c, 6c**), even though cell nuclei in the cortical layers of *SHANK3*- /-, but not *SHANK3*+/-, organoids were significantly smaller than those in iCtrl organoids (**Supplementary Fig. 14**).

**Figure 6.**
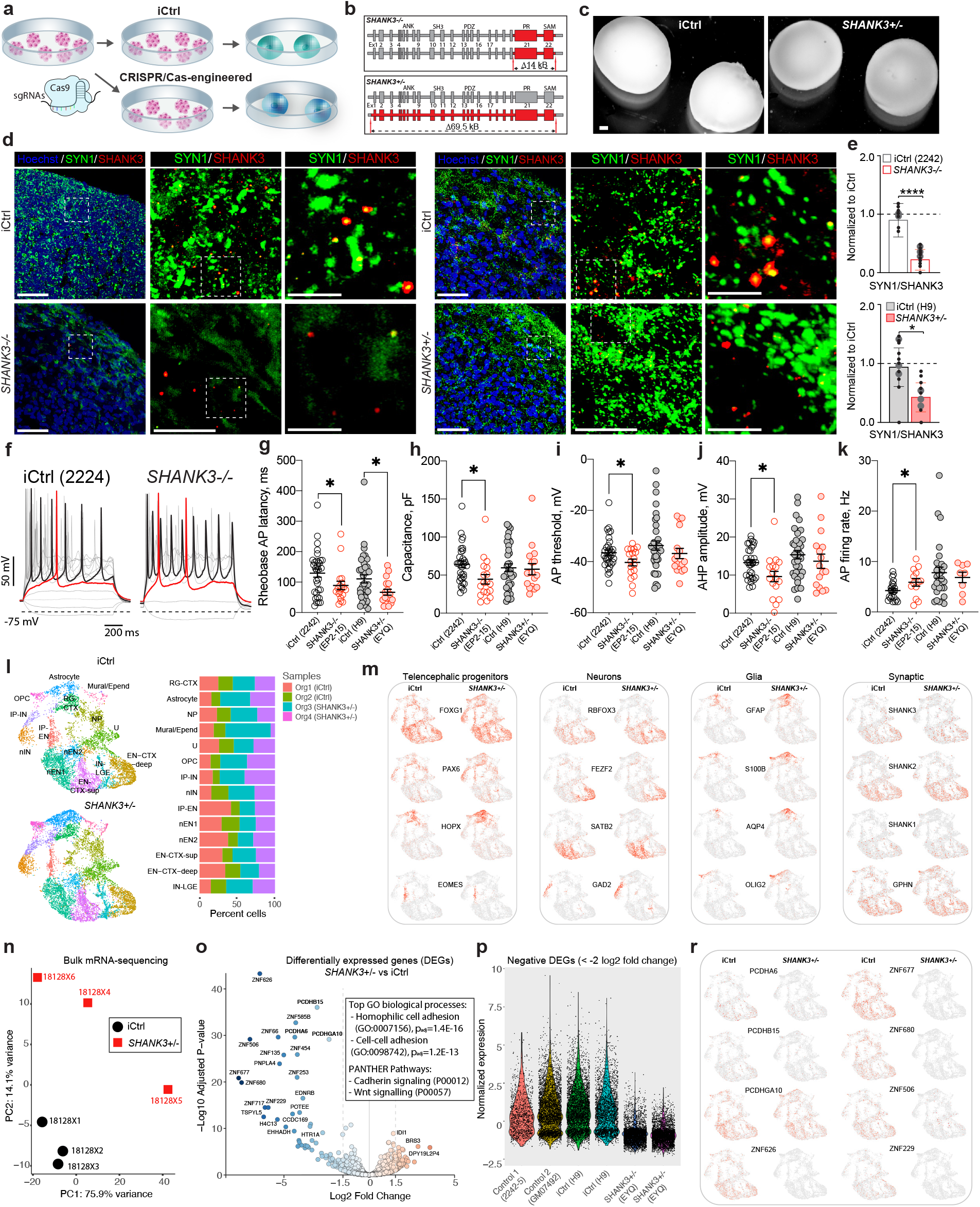
Cellular and molecular functional deficits detected in SHANK3-deficient SNR-derived organoids. **a-b**, Generation of CRISPR/Cas9-engineered PSC lines and organoids with homozygous and complete hemizygous SHANK3 deletions (*SHANK3-/-* and *SHANK3+/-*, respectively). **c**, Images of 5-month-old iCtrl and *SHANK3+/-* SNR-derived organoids. **d**, Synaptic stainings using anti-SYN1 and anti-SHANK3 antibodies in iCtrl and SHANK-deficient organoids (left: iCtrl [2242-5] and *SHANK3-/-* [EP2-15]; right: iCtrl [H9] and *SHANK3+/-* [EYQ]). **e**, Quantification of SYN1/SHANK3 puncta (top: n = 7 and 12 sections from 2 iCtrl [2242-5] and 3 *SHANK3-/-* [EP2-15] organoids, respectively; bottom: n = 11 and 11 sections from 3 iCtrl [H9] and 3 *SHANK3+/-*[EYQ] organoids, respectively). **f**, Traces of voltage deflections recorded from iCtrl (2242-5) and *SHANK3-/-* (EP2-15) neurons in response to different somatic current injections (ΔI=10pA). **g-j**, Quantified Rheobase AP latency (**g**), membrane capacitance (**h**), AP threshold (**i**), AHP (**j**), and AP firing rate (**k**) for iCtrl and SHANK3-deficient neurons (n = 32, 18, 37, and 18 cells, from 4 iCtrl [2242-5], 3 *SHANK3-/-* [EP2-15], 11 iCtrl [H9], and 2 *SHANK3+/-* [EYQ] organoids) in 5-month-old SNR-derived organoid slices. **k**, UMAP visualizations of unsupervised clustering of scRNA-seq data from 2 iCtrl [H9] and 2 *SHANK3+/-* [EYQ] SNR-derived organoids and relative cluster composition (right). These organoids are included in the clustering results presented in Fig. 3. **l**, Heatmap of expression of different cell-type specific marker genes in iCtrl (H9) and *SHANK3+/-* (EYQ) organoids. **m**, PCA plot of bulk RNA-seq data from 3 iCtrl (H9) and 3 *SHANK3+/-* (EYQ) organoids. **n**, Volcano plot visualization of bulk RNA-seq data and top GO and PANTHER pathways. **o**, Expression of bulk RNA-seq DEGs in cells from 3 Ctrl and 2 *SHANK3+/-* organoids. **p**, Normalized expression of bulk RNA-seq DEGs in scRNA-seq datasets. **r**, scRNA-seq heatmaps of the expression of protocadherins and zinc-finger genes that are downregulated in *SHANK3+/-* organoids. Data are mean ± s.e.m.; *P<0.05, ****P<0.001, unpaired t-test. Scale bars = 450 (**c**), and 50, 20, and 5 μm (**d**).

Excitatory synaptic deficits and elevated intrinsic excitability have been observed in association with SHANK3 deficiency in both human and rodent neurons (Chiola et al., 2021; Jiang and Ehlers, 2013; Peixoto et al., 2016; Shcheglovitov et al., 2013; Yi et al., 2016). Therefore, we investigated the numbers of excitatory synapses in 5-month-old SHANK3-deficient organoids using synaptic immunostaining (**Fig. 6d-e**) and the intrinsic excitability using slice patch-clamp electrophysiology (**Fig. 6f-k, Supplementary Fig. 15**, and **Supplementary Table 4**). We found significantly reduced numbers of Synapsin1/SHANK3-containig excitatory synaptic puncta in both *SHANK3-/-* and *SHANK3+/-* organoids as compared to the respective iCtrl organoids (**Fig. 6e**). We also observed that SHANK3-deficient neurons were more excitable than iCtrl neurons (**Fig. 6f-k**), exhibiting a shorter AP latency in both *SHANK3-/-* and *SHANK3+/-* neurons (**Fig. 6g**) and smaller membrane capacitance (**Fig. 6h**), more hyperpolarized AP threshold (**Fig. 6i**), reduced amplitude of after-hyperpolarization potential (AHP) (**Fig. 6j**), and increased firing rate (**Fig. 6k**) in *SHANK3-/-* neurons. These results are consistent with the idea that SHANK3 is an important regulator of intrinsic neuronal excitability and that the level of SHANK3 expression influences the severity and/or manifestation of the cellular deficits in human neurons.

To gain insights into the molecular pathways disrupted by hemizygous *SHANK3* deletion in SNR-derived organoids, we analyzed a subset of the 5-month-old scRNA-seq data (**Fig. 3e**) that includes only samples obtained from two pairs of iCtrl and *SHANK3*+/- organoids (**Fig. 6l**) and performed bulk RNA sequencing on additional three pairs of 5-month-old iCtrl and *SHANK3*+/- organoids (**Fig. 6n**). Although we observed no major differences in the cellular composition between iCtrl and *SHANK3*+/- organoids based on the expression pattern of different cells type specific markers (**Fig. 6l-m**), *SHANK3*+/- organoids showed significantly reduced expression of several clustered protocadherins, including *PCDHA6, PCDHGA10*, and *PCDHB15*, and multiple primate-specific zinc finger (ZNF) genes, including *ZNF626, ZNF677, ZNF680, ZNF506*, and *ZNF229* using bulk RNA-seq (**Fig. 6o** and **Supplementary Table 5**). Consistent with reduced expression of multiple clustered protocarderins in *SHANK3*+/- organoids, the gene ontology (GO) analysis using the PANTHER platform (Mi et al., 2021) identified the cadherin (P00057) and Wnt (P00057) signaling pathways as significantly overrepresented (**Fig. 6o** and **Supplementary Table 6**). We confirmed that the most differentially expressed genes (DEGs) identified using bulk RNA-seq were significantly downregulated in scRNA-seq datasets obtained from *SHANK3*+/- organoids as compared to isogenic and unrelated control organoids (**Fig. 6p-r** and **Supplementary Fig. 16)**. Interestingly, we observed no strong enrichment of the bulk RNA-seq DEGs in specific neuronal clusters, suggesting that multiple cell types could be affected by SHANK3 deficiency in the developing human telencephalon.

## DISCUSSION

Human stem cell-derived organoids have provided unique opportunities to produce organized, region-specific brain tissue *in vitro* to model the early aspects of human brain development and disorders (Eiraku et al., 2008; Kadoshima et al., 2013; Lancaster et al., 2013; Mariani et al., 2012; Paşca et al., 2015; Qian et al., 2016; Sloan et al., 2017; Watanabe et al., 2017). However, organoid-to-organoids variability, unpredictable cytoarchitecture, slow developmental maturation, and abnormal cell-type specification have been the major issues limiting the overall enthusiasm of organoid models (Bhaduri et al., 2020; Quadrato et al., 2017). The present study describes a new method for generating human cortico-striatal organoids from stem cell-derived SNRs. We comprehensively characterized the properties and organizations of different cell types in these organoids and obtained novel insights into the molecular programs associated with their specifications. We also demonstrated that 5-month-old SNR-derived organoids contain neurons and neural networks that show signs of functional and morphological maturity, including repetitive APs, excitatory and inhibitory synaptic currents, elaborate dendritic arbors and spines, and functional neural networks. Finally, we used SNR-derived organoids to investigate the cellular and molecular deficits caused by homo- and hemizygous *SHANK3* deficiencies. Homozygous SHANK3-deficient organoids had multiple neurodevelopmental deficits associated with impaired neurogenesis, morphogenesis, synaptogenesis, and intrinsic excitability; while only reduced numbers of excitatory synapses and slightly elevated intrinsic excitability were detected in organoids with *SHANK3* hemizygosity. Interestingly, *SHANK3* hemizygosity caused reduced expression of multiple clustered protocadherins and primate-specific ZNF genes in SNR-derived organoids. In summary, this reveals novel molecular deficits caused by *SHANK3* deficiencies in human cortico-striatal tissue and validates SNR-derived organoids as a reliable model for studying human telencephalic development under normal and pathological conditions.

### Using SNRs to generate organoids

The use of SNRs for organoid formation offers several potential advantages. First, it mimics the development of neuroectodermal tissue from a singular neural tube *in vivo*. Previous studies characterized organoids generated from pluripotent or neural stem cells that contained multiple neural rosettes (Birey et al., 2017; Eiraku et al., 2008; Kadoshima et al., 2013; Lancaster et al., 2013; Mariani et al., 2012, 2015; Paşca et al., 2015; Qian et al., 2016; Quadrato et al., 2017). The presence of multiple rosettes per organoid can result in an unpredictable cytoarchitectural and neuronal organization. We observed that NPs and neurons in SNR-derived organoids are predictably organized around the lumen, which allowed us to find the functionally mature neurons with relatively high efficiency. In addition, neural tissue with multiple rosette-like structures has been identified as a hallmark of several types of brain-related tumors (Louis P. Dehner, 1993; Werbowetski-Ogilvie et al., 2012; Wippold and Perry, 2006), suggesting that multiple neural rosettes in an organoid may contribute to the development of unanticipated abnormalities that could confound disease modeling studies. Organoid generation from SNRs overcomes these limitations.

Despite several important advantages, our approach has some limitations. First, although manual isolation of SNRs is an effective approach for generating organoids with a single lumen, some (up to 50%) of SNR-derived organoids may still contain multiple lumens or without lumens at all due to errors in SNR selection. To overcome this limitation, only SNR-derived organoids with a clearly visible single lumen were embedded in Matrigel. Another limitation may be associated with the use of Matrigel as a scaffold. The Matrigel composition is not well-defined and may vary batch-to-batch (Hughes et al., 2010). Matrigel may also cause local disruptions in organoid organization by irregular gelling. Future experiments should be directed towards the development of novel well-defined biomaterials to substitute for Matrigel.

### LGE-derived inhibitory neurons in SNR-derived organoids

Although the human cortex and striatum contain an increased number and greater diversity of INs (Krienen et al., 2020; Rakic, 2009; Yuste, 2005), our understanding of the origins and underlying molecular programs remains limited. Most previous studies have been focused on the investigation of MGE-derived NPs as the major source of INs in the cortex and striatum (Anderson et al., 1997; Hu et al., 2017). We demonstrate that human cortico-striatal SNR-derived organoids contain no detectable MGE-like NPs (NKX2-1- and LXH6-expressing cells), and the INs in organoids are produced from LGE-like progenitors expressing *GSX2, SOX6, ASCL1, DLX1* and *DLX2*. In the mouse brain, LGE-derived NPs predominantly contribute to the production of GABAergic primary striatal neurons and interneurons that migrate to the olfactory bulb (Stenman et al., 2003). In the human and primate brains, LGE-derived NPs have also been implicated in the production of cortical interneurons (Ma et al., 2013). The expression of typical cortical and striatal GABAergic interneuron markers, such as CALR, CALB, SST, and PV, in SNR-derived organoids suggest the cortico-striatal identities of these cells. Importantly, we observed that at least a fraction of INs in SNR-derived organoids are synaptically integrated into organoid networks and may contribute to their maturation.

Striatal projection neurons were recently produced in organoids using an agonist of retinoid X receptors, SR11237 (Miura et al., 2020). In the present study, we achieved the generation of striatal neurons by using vitamin A-containing differentiation media for neural induction and differentiation. Vitamin A (or retinol) is the immediate precursor of retinoic acid, an important regulator of both cortical and striatal development (Chatzi et al., 2011; Rataj-Baniowska et al., 2015; Shi et al., 2012; Siegenthaler et al., 2009). It remains to be determined whether striatal projection neurons generated in SNR-derived organoids are functionally mature and capable of establishing functional unidirectional connections with cortical neurons (Bolam et al., 2000).

### Ependymal cells, oligodendrocytes, and mural cells in SNR-derived organoids

We showed that 5-month-old SNR-derived organoids contain a diverse array of non-neuronal cells, including astrocytes, oligodendrocytes, pericytes, and ependymal-like cells. Although astrocytes and oligodendrocytes have previously been reported in organoids (Madhavan et al., 2018; Marton et al., 2019), the detection of maturing MBP-expressing oligodendrocytes in 5-month-old SNR-derived organoids was unexpected. The generation and maturation of cortical oligodendrocytes correlates with the level of functional neuronal maturation (Barres and Raff, 1993; Stevens et al., 2002). Thus, the presence of oligodendrocytes in 5-month-old SNR-derived organoids may be associated with improved functional maturation of neurons and neural networks. However, additional experiments will be necessary to fully characterize the properties of OPC and oligodendrocytes in SNR-derived organoids.

Brain pericytes are essential for the establishment of the blood-brain barrier (Daneman et al., 2010). Dysfunctional pericytes have been implicated in neurodegenerative disorders, stroke, and neuroinflammation (Rustenhoven et al., 2017). However, the developmental origin of brain pericytes is debatable (Korn et al., 2002). Our results are consistent with the idea of the neuroectodermal origin of pericytes in SNR-derived organoids, as we generated organoids from isolated SNRs and found no evidence of the presence of mesodermal cells in SNR-derived organoids using scRNA-seq.

We were intrigued to find FOXJ1-expressing cells lining the lumen in 5-month-old SNR-derived organoids. FOXJ1 is a relatively specific marker for ependymal cells, which are ciliated glial cells that line the ventricles in the developed brain. Although no cilia were observed on the FOXJ1-expressing cells, scRNA-seq identified increased expression of genes important for ciliogenesis, including *FOXJ1, RFX2*, and *ZMYND10* (Thomas et al., 2010; Zariwala et al., 2013).

We propose that SNR-derived organoids may be useful for studying glial and mural cell development under normal and pathological conditions.

### Functional maturation of neurons and neural networks in SNR-derived organoids

In our experiments, 86% (103/120) of recorded cells in 5-month-old SNR-derived organoids were functionally active and could be subdivided into five groups based on the functional properties. Neurons in different groups exhibited different passive membrane properties such as input resistance, resting membrane potential, capacitance, and AP firing profiles, which could be related to different developmental status or neuronal identity. However, additional experiments will be needed to determine the developmental status and identities of neurons in different clusters and to compare them with neurons in the mouse or human brain.

We found that about 50% recorded neurons were synaptically active. This is comparable to what was observed in the previous studies on organoids (Birey et al., 2017; Qian et al., 2016). Consistent with the synaptic maturation of neurons in SNR-derived organoids, we observed spontaneous electrical activity and functional neural networks capable of generating oscillatory rhythms in 6-month-old SNR-derived organoids. This is relatively early as compared to a previous study on cerebral organoids (Quadrato et al., 2017), in which spontaneous activity was observed only after 8 months in culture. However, our results are comparable to those observed in organoids plated on 2D multi-electrode arrays (Trujillo et al., 2019). While the cellular and circuit mechanisms responsible for the generation of rhythmic electrical activity in human neural networks remain unknown, it is believed that low-frequency oscillations emerge from long-range connections while high-frequency oscillations emerge from shorter ones (Uhlhaas and Singer, 2012). Detection of both low-(4-30 Hz) and mid- to high-frequency (30-100 Hz) oscillations in SNR-derived organoids may indicate the presence of both long- and short-range functional connections. In addition, high-frequency oscillations rely on the activity GABAergic interneurons (Cardin et al., 2009), and we confirmed the presence of different subtypes of functional GABAergic interneurons in SNR-derived organoids. Interestingly, even stronger oscillations (> 100 Hz) have been detected in organoids containing MGE-derived inhibitory neurons (Samarasinghe et. al., 2019; Trujillo et al., 2019). This is consistent with the idea that PV-expressing MGE-derived inhibitory interneurons are essential for generation of high-frequency oscillations in cortical neural networks (Sohal et. al., 2009). More experiments will be required to elucidate the contributions of different subtypes of inhibitory interneurons to the development of synchronized oscillations in organoids.

Our approach for measuring electrical activity in organoids could be useful for studying the cellular and molecular mechanisms involved in functional neural network establishment under normal and pathological conditions and for investigating the effects of neural stimulation (i.e., electrical, magnetic, ultra/infrasonic, and infrared) on human neural cells and networks.

### Cellular and molecular deficits in SHANK3-deficient SNR-derived organoids

Mutations and deletions in *SHANK3* have been identified in most patients with Phelan-McDermid syndrome (Harony-Nicolas et al., 2015; Mitz et al., 2018; Soorya et al., 2013) and approximately 2% of individuals with idiopathic autism and intellectual disability (Leblond et al., 2014; De Rubeis et al., 2014). SHANK3 deficiency was shown to cause excitatory synaptic and intrinsic excitability in both rodent and human neurons (Chiola et al., 2021; Jiang and Ehlers, 2013; Peixoto et al., 2016; Shcheglovitov et al., 2013; Yi et al., 2016), but the underlying molecular mechanisms remain largely unknown. In this study, we confirmed the presence of synaptic and intrinsic deficits in SHANK3-deficient organoids and identified disrupted signaling cascades that may contribute to SHANK3-mediated deficits. Specifically, *SHANK3*+/- organoids had disrupted expression of multiple primate-specific ZNF genes (Huntley et al., 2006) and several clustered protocadherins. Many ZNF genes have been recently shown to be involved in human brain development (Al-Naama et al., 2020), but their roles remain poorly understood. Clustered protocadherins, similar to *SHANK3*, have been implicated in controlling neural circuit assembly, dendritic morphogenesis, and spinogenesis (Canzio and Maniatis, 2019; Lefebvre et al., 2012; Suo et al., 2012). In addition, mutations in protocadherins have been detected in individuals with autism (Anitha et al., 2013), and disrupted protocadherin expression has been observed in stem cell-derived neurons from individuals with major depression (Vadodaria et al., 2019) and schizophrenia (Shao et al., 2019). Future studies should determine how loss of *SHANK3* causes reduced expression of ZNF genes and cluster protocadherins and whether targeting of specific genes could compensate for deficits caused by SHANK3 deficiency.

## Materials and Methods

### Stem cell culture and neural differentiation

Human pluripotent stem cell lines used in this study (control lines: H9 [WiCell], 2242-5 [Dolmetsch lab, Stanford University], and GM07492 [Ernst lab, McGill University (Bell et al., 2018)]; SHANK3-defient lines: EYQ2-20 [SHANK3+/-, Shcheglovitov lab, (Chiola at al., in press)] and EP2-15 [SHANK3-/-, Shcheglovitov lab]) were maintained on Matrigel (1%, BD) in Essential 8 Medium (Thermo Fisher Scientific). For neural differentiation, cells were passaged at high density to reach ∼90% confluency 2-3 days post-passaging. For induction, E8 medium was replaced with neural differentiation (ND) medium containing a 1:1 mixture of N2 medium (DMEM/F-12, 1% N2 Supplement [Cat#17502048], 1% MEM-NEAA, 2 µg/ml Heparin, and 1% Pen/Strep) and B-27 medium (Neurobasal-A [Cat#10888022], 2% B27 Supplement with vitamin A [Cat#17504044], 1% GlutaMAX and 1% Pen/Strep) supplemented with SMAD inhibitors (4 µM dorsomorphin and 10 µM SB431542). ND medium was replaced daily for the next 7–10 days. On days 7–10, cells were manually scraped from the plate using a sterile glass hook. Cell clumps were re-plated on Matrigel-coated plates in ND medium with EGF (10 ng/mL) and FGF (10 ng/mL). Approximately 50–100% of media was replenished daily for the next 3–6 days to remove dying cells. Neural rosette clusters appeared after ∼6 days in EGF/FGF-containing media and were manually split using a sterile glass hook and re-plated on Matrigel-coated plates at low density. On days 15–19, SNRs were manually isolated and transferred to ultra-low attachment plates in ND medium with EGF (10 ng/mL) and FGF (10 ng/mL). Plates were kept on an orbital shaker (45 rpm) in a 37°C, 5% CO_2_ incubator with 50% media exchange every other day to maintain the final concentrations of EGF and FGF at 10 ng/mL. At 2–3 weeks post SNR isolation, SNR-derived organoids with clearly visible single lumen were transferred into Matrigel as previously described (Lancaster and Knoblich, 2014) and cultured in uncoated 10-cm Petri dishes with ND medium without growth factors. The dishes with organoids were transferred to the orbital shaker 24 h post-Matrigel embedding. Approximately 50% of the media was exchanged every 3–4 days thereafter. At 3 months post SNR isolation, BDNF (10 ng/mL), GDNF (10 ng/mL), and NT-3 (10 ng/mL) were added to the culture with each media change to promote synapse development and maturation.

### Generation of isogenic stem cell line with homozygous *SHANK3* deletion

A *SHANK3*-deficient stem cell line (EP2-15) was generated using a CRISPR/Cas9 approach from the 2242-5 iPSC line. Two sgRNA sequences, gRNA #8 (TGAGCCGCTATGACGCTTCAGGG) and gRNA #4 (TTTCTCAGGGGTCCCCGGTGGGG), designed to flank exons 21-22 and the stop codon for creating an ∼14-kB deletion, were introduced using electroporation together with Cas9 sequence under the control of the EF-lα promoter. Upon electroporation of 5 µM CRISPR/Cas9 plasmids into ∼3×10^6^ cells using the Amaxa Nucleofector System (program A023), iPSCs were resuspended in E8 medium with ROCK inhibitor. The following day, cells were exposed to 0.25 µg/mL puromycin for 48 h. Antibiotic-resistant colonies were picked and expanded on Matrigel-coated plates in E8 medium. A total of 82 single colonies have been manually picked, propagated, and screened for the deletion using conventional PCR and a set of one forward and two reverse primers (Primer F [yw3]: CTCAGGGCCTGCTTGATGAC; Primer R [yw4]: GTGCGCTCCTGAAGGACAAT; and Primer R [yw7]: GGTCTTGCATCGAGGTGCTC), which recognize different regions in the proximity of exons 21-22 (**Fig. 6a**). Out of the 82 colonies, one was identified with homozygous *SHANK3* deletion encompassing exons 21-22. The deletion was verified using PCR product purification, subcloning into a TOPO TA cloning vector (Thermo Fisher), and sequencing.

### Single-cell RNA-sequencing

#### Single-cell collection from 1-month-old organoids

SNR-derived organoids were rinsed with sterile Dulbecco’s phosphate-buffered saline (DPBS, 1X) containing 5% DNaseI and then dissociated in a pre-warmed working solution of either Accutase (1 month) or papain (5 month) with 5% DNase I. The samples were incubated at 37°C for 5 min and centrifuged for 1 min at 1000*g*. After removal of the enzyme supernatant, cells were resuspended in Trypsin Inhibitor with 5% DNase I, incubated for 5 min at 37°C, and centrifuged for 1 min at 1000*g*. Cells were washed with Neurobasal-A Medium with 5% DNaseI and briefly centrifuged before the next step. Cells were resuspended in 200 µL Neurobasal-A and manually triturated by pipetting up and down approximately 20–30 times. The suspension was centrifuged at 1600 rpm for 4 min and resuspended in 200– 500 µL Neurobasal-A. The suspension was passed through a 40-µm cell strainer to yield a uniform single-cell suspension. Samples were kept on ice from this point forward. Samples were centrifuged at 1200–1500 rpm for 6–8 min. After removal of the supernatant, cells were resuspended in 35–50 µL Hank’s Balanced Salt Solution (1X) or DPBS (1X), and cell viability was assessed using Trypan blue staining. Samples were quickly transferred to the High Throughput Genomics Core Facility at the University of Utah.

#### Single-cell collection from 5-months-old organoids

SNR-derived organoids were subjected to vibratome slicing in ice-cold “cutting” artificial cerebrospinal fluid (ACSF), containing: KCl 2.5 mM, MgCl_2_ 7 mM, NaH_2_PO_4_ 1.25 mM, Choline-Cl 105 mM, NaHCO_3_ 25 mM, D-Glucose 25 mM, Na^+^-Pyruvate 3 mM, Na^+^-L-Ascorbate 11 mM, CaCl_2_ 0.5 mM. Collected tissue slices (350 µm thick) were enzymatically dissociated using prewarmed Papain-containing solution for 20 to 30 minutes at 37°C. After removing enzyme supernatant, cells were resuspended in Trypsin Inhibitor with 5% DNase I, incubated for 10 minutes at 37°C, and gently triturated with p1000, p200 and p20 in the trypsin inhibitor solution. The cell suspension was passed through a 40 µm cell strainer. Samples were centrifuged at 1000 rpm for 7 minutes to remove supernatant. After removing supernatant, cells were resuspended in ice-cold sterile PBS (1X) with 0.04% bovine serum albumin (BSA) to 1,000 cells/µL. Samples were transferred on ice to the High Throughput Genomics Core Facility at the University of Utah to assess cell viability using Trypan Blue staining and to load the cells in the 10X Genomics Chromium Single Cell 3′ Microfluidics Chip (10,000 cell/channel). Using this approach, we achieved more than 90% cell viability.

#### Single-cell Capture and Library Preparation

Single-cell capture, lysis, and cDNA synthesis were performed with the 10x Genomics Chromium at the High Throughput Genomics Core Facility at the University of Utah. All cells were processed according to 10X Genomics Chromium Single Cell 3′ Reagent Guidelines. Sequencing was performed using Illumina HiSeq 26 x 100 cycle paired-end sequencing.

#### Single-cell RNA-seq processing

Reads were processed using the 10X Genomics Cell Ranger 2.0 pipeline as previously described (Zheng et al., 2017) and aligned to the human GRCh38 reference assembly (Ensembl v84). Gene-barcode matrices were constructed for each sample by counting unique molecular identifiers (UMIs). The single-cell RNA-seq data reported in this paper is available in the GEO repository with accession number GSE118697.

### Single-cell RNA-seq analysis of 1-month-old organoids

#### Data filtering

To exclude low quality cell libraries, we performed quality control using the scater R package (McCarthy et al., 2017) and filtered cells based on the distribution of library size and mitochondrial RNA. Cells with less than 2000 UMIs or 1200 detected genes and those with more than 14000 UMIs or 4000 detected genes were removed. To remove cells that contain abnormally high or low number of UMIs relative to its number of detected genes, we fit a smooth curve (loess, span= 0.5) to the number of UMIs vs. the number of genes after log-transform, and removed cells with residuals more than three standard deviations from the mean. We also removed cells with more than 6% of UMIs assigned to mitochondrial genes. After cell filtering, genes detected with at least one UMI in at least two cells were kept in the dataset.

In addition, we used the DoubletDetection (Version v2.4) Python package (https://zenodo.org/record/2678042) to remove putative doublet or multiplet cell libraries from the dataset. We run the program for 50 iterations for each sample separately, we used a conservative threshold and classified cells with p < 10^−5^ in 90% of iterations as multiplets. Using this method, 2.5% of cells were removed, which agrees with the expected multiplet rate of the 10X Genomics platform (2.3% for ∼3000 cells recovered). The filtered dataset containing 9577 cells were used for downstream analysis. We performed library size normalization, feature selection, dimensionality reduction and clustering using the Seurat R package (Butler et al., 2018).

#### Data normalization

The gene expression of each cell was normalized by its total UMI count with a default scale factor of 10000, and log-transformed the normalized count for downstream analysis.

#### Feature selection and scaling

We used the FindVariableGenes function in Seurat to identify a set of highly variable genes (HVGs), which are informative genes that exhibit high variance across the dataset. We reason that the gene expression values in each single cell library are determined by the combination of four variables: (1) biological variables related to cell type identity, (2) biological variables related to common cellular activities rather than a specific cell type (e.g. cell cycle), (3) technical variables related to library size (e.g. number of UMIs), and (4) random noise. To focus our downstream analysis on gene expression programs involved in cell type identity, we performed careful selection of HVGs to eliminate the effects unrelated to cell type. To exclude genes with very low expression and housekeeping genes from HVGs, we only considered genes that are detected in at least 10 cells but less than 90% of all cells, with average expression between 0.03 and 3. Among these genes, the top 1200 genes ranked by scaled dispersion were selected as HVGs. We further excluded ribosome and mitochondrial gene (as identified by gene names starting with RPL, RPS or MT) from HVGs.

To remove the effect of cell cycle signals on dimensionality reduction and cell type clustering, we used the list of 97 cell cycle genes provide by the Seurat package as the set of known cell cycle genes. Using this information, we first removed 53 known cell cycle genes present in the initial set of HVGs. Then we used these genes as reference to identify other putative cell cycle genes in HVGs by correlation analysis: (1) we collected a subset of robust cell cycle genes that show strong correlation (spearman correlation > 0.3) with any other known cell cycle genes; (2) next, we identified another 88 genes in HVGs that are highly correlated (spearman correlation > 0.3) with any cell cycle gene in the above subset. These putative cell cycle genes were subsequently removed from HVGs. The final set of HVGs consisted of 1056 genes.

To remove the sample-specific batch effect, we performed batch correction using the mutual nearest-neighbors correction method in the scran R package (Haghverdi et al., 2018). Batch correction was performed on HVGs with the sigma value of 0.5. The batch-corrected HVG expression matrix was then fed into the ScaleData function in Seurat. The ScaleData function removes the effect of library size on cell-cell variation in gene expression by fitting a linear regression model on each gene using the number of detected UMIs as a predictor. The residuals were then scaled to zero-mean and unit-variance.

#### Dimensionality reduction

To reduce the dimensionality of the dataset, we performed principal component analysis (PCA) on the scaled HVG expression matrix. The first 21 principal components were chosen for downstream analysis, as judged by the PCA elbow plot and the heatmap distribution of top-ranked genes for each principal component.

#### Cell type clustering

Clustering of single cells were performed using the FindClusters function in Seurat. Briefly, we built a shared nearest neighbor graph of single cells using the first 21 principal components and a neighborhood size of 30. Louvain clustering with resolution of 1.2 was then performed on the single cell graph to partition cells into discrete clusters. We computed a cosine similarity matrix between all pairs of clusters using cluster-averaged expression profile and constructed a cluster dendrogram using (1 - cosine_similarity) as the distance metric and the average linkage method.

#### Differential expression

To find marker genes that are enriched in each cluster, we performed differential expression analysis between each cluster and the rest of the dataset using the Wilcoxon test in the Seurat FindAllMarkers function. We require marker genes to be present in at least 25% of cells in the cluster, have a fold change of 2 by average expression level, a fold change of 2 by the percentage of expressing cells, and a Bonferroni-corrected p value < 0.01.

#### Nonlinear dimensionality reduction

To embed single cells into a 2-dimensional visualization, we took the above principal components as inputs and performed nonlinear dimensionality reduction using the Uniform Manifold Approximation and Projection (UMAP) algorithm (McInnes and Healy, 2018) with n_neighbors = 20 and min_dist = 0.3. The resulting UMAP embedding attempts to preserve both the local neighbor relations and the global structure of the data.

#### RNA velocity analysis

RNA velocity analysis was performed using the Velocyto Python package (La Manno et al., 2018) to infer the future states of single cells during development. Briefly, RNA velocity vectors for each cell were calculated using the relative abundance of spliced and unspliced mRNAs of each gene. Spliced and unpliced read counts were generated using the Velocyto command line interface. Genes with low spliced counts (less than 40 UMIs or detected in less than 30 cells) were excluded from analysis. The top 3000 variable genes were selected by fitting the variance vs. mean relationship.1648 variable genes are kept after filtering by unspliced counts (less than 25 UMIs or detected in less than 20 cells), cluster-averaged expression (unspliced < 0.01 or spliced < 0.08 in all clusters), as well as removing ribosome, mitochondria and cell cycle genes. PCA was performed on variable genes and the first 20 principal components were used to build a k-nearest neighbor graph (KNN) with k = 200. The velocity vector was inferred after KNN imputation and the future state of each cell was extrapolated using the constant velocity model and a time step of 1. To project the velocity vectors onto the single cell UMAP embedding, the transition probabilities between pairs of cells were estimated with a neighborhood size of 750 and the embedding shift was calculated with sigma = 0.05. The smoothed velocity vector grid was calculated with a smoothing factor of 0.8 and neighborhood size of 300. To simulate the progression of single cells through developmental time, Markov Chain simulation was performed for 200-time steps.

#### Smoothed gradient of gene expression change using Slingshot

As UMAP constructs a low-dimensional embedding of the data manifold that captures both local and global structure of the data, we fit smooth curves along the UMAP embedding to study how gene expression changes across cell types. The Slingshot algorithm (Street et al., 2018) was used to fit both linear and branching curves across the single cell clusters. To constrain the smooth curve fitting algorithms to recover biologically meaningful solutions, we (1) performed the fit separately on the progenitor clusters, the excitatory lineage and the inhibitory lineage, and (2) used the Markov Chain endpoint clusters from RNA velocity inference as the supervised end points in the Slingshot algorithm. This method recovered smooth curve trajectories that agrees well with the RNA velocity analysis. Cells along each lineage were projected onto the Slingshot trajectory curves and ordered according to the assigned pseudotime values. Generalized additive model from the gam R package was used to identify genes that are differentially expressed along each trajectory lineage. Genes with small fold change (less than 6 for neural differentiation trajectories or less than 3 for the progenitor trajectories) were removed from the list of differentially expressed genes. Genes were assigned to early- or late-stage by the time of their peak expression along the trajectory. A set of common early EN/IN differentiation genes were produced by the intersection of early EN/IN genes across neural differentiation trajectories.

#### Gene Ontology (GO) analysis

GO analysis was performed using the ToppGene Suite (Chen et al., 2009).

### Single-cell RNA-seq analysis of 5-month-old organoids

Single-cell RNA-seq analysis of the six 5-month-old organoids was performed using the similar methods as the 1-month-old organoids.

#### Data filtering and normalization

To exclude low quality cell libraries, we filtered cells based on the distribution of library size and mitochondrial RNA. In Org5, cells with less than 2000 UMIs or 750 detected genes and those with more than 23000 UMIs or 3800 detected genes were removed. For the other five organoids, cells with less than 1300∼1500 UMIs or 700 detected genes and those with more than 35000 UMIs or 7000 detected genes were removed. To remove cells that contain abnormally high or low number of UMIs relative to its number of detected genes, we fit a smooth curve (loess, span= 0.5) to the number of UMIs vs. the number of genes after log-transform, and removed cells with residuals more than three standard deviations from the mean. We removed cells with more than 6∼10% of UMIs assigned to mitochondrial genes. After cell filtering, genes detected with at least one UMI in at least two cells were kept in the dataset. We performed multiplet detection using DoubletFinder (McGinnis et al., 2019) in organoid samples with more than 1500 cells and removed the putative multiplets. The preprocessed individual datasets contain 4682, 2823, 4961, 4618, 1152, 4250 cells, respectively.

#### Feature selection and scaling

We selected 3000 HVGs separately for each of the six 5-month-old organoids using the SCTransform function in Seurat v3 (Hafemeister and Satija, 2019; Stuart et al., 2019). SCTransform also normalized and scaled the expression matrix. Cell cycle genes were then removed from HVGs following the same approach as in the 1-month-old organoids.

#### Integration of the 5-month-old organoid datasets

To perform integrated analysis on the six 5-month-old datasets, we combined them together using the single-cell integration pipeline in Seurat v3. The set of gene features used is the union of HVGs from all datasets. To ensure all features are present in each dataset, we padded zeros rows when a gene within the union features is absent in a dataset’s gene expression matrix. We selected 3000 integration features using the SelectIntegrationFeatures function, and prepared the dataset for integration using the PrepSCTIntegration function. A set of “anchors” were identified between pairs of datasets using the FindIntegrationAnchors function (dims = 1:30, k.anchor = 3, k.filter = 20, k.score = 15). Then, we integrated all datasets together using these anchors and the IntegrateData function (dims = 1:30, k.weight = 50). Clustering was performed on the integrated dataset using a resolution of 0.55, a neighborhood size of 20 and the first 30 principle components. UMAP was performed with n_neighbors = 20 and min_dist = 0.3. Subsequent differential expression analysis and marker gene selection was done using the same method as in the 1-month-old organoids. To analyze the correlation of cell type clusters between a pair of different datasets, we computed the cluster-averaged gene expression profile for each cluster using the intersection of HVGs from the two datasets and the scaled expression values. Pearson correlation was then performed between clusters from the two datasets.

### Bulk RNA-seq and correlation analysis

Samples from 5-month-old SNR-derived organoids were homogenized in Trizol (Invitrogen), and total RNA was purified using the RNeasy Micro Kit (Qiagen) with DNase I digestion. Ribosomal RNA was depleted with the NEBNext rRNA Depletion Kit. Libraries were prepared using the NEBNext Ultra Directional RNA Library Prep Kit for Illumina and verified using the Bioanalyzer RNA Nano Assay (Agilent) before sequencing on the NextSeq 500 (Illumina). Single-end reads were trimmed to 70 bp for genomic alignment. Total RNA-seq reads for SNR-derived organoids and fetal human brain samples were aligned to hg38 (i.e., GRCh38.p2) using BWA and further processed for exonic expression levels (RPKM) across ∼ 24,000 RefSeq-annotated genes using a previously described pipeline (Ataman et al., 2016). Reads that did not map uniquely with at most two mismatches were excluded.

The correlation analysis was performed on data from 7 SNR-derived organoids and 10 fetal-brain samples (Ataman et al., 2016), plus 15 samples from 4 published studies available through the NCBI Gene Expression Omnibus (GEO) site: GSM2580319, GSM2580321, GSM2580323, GSM2580325, GSM2580327, and GSM2580329 (Watanabe et al., 2017); GSM2112671 and GSM2112672 (Qian et al., 2016); GSM3408648, GSM3408667, and GSM3408685 (Yoon et al., 2019); and GSM2180144, GSM2180145, GSM2180138, and GSM2180139 (Luo et al., 2016). Twelve datasets of FASTQ reads (GSM2580319, GSM2580321, GSM2580323, GSM2580325, GSM2580327, GSM2580329, GSM2112671, GSM2112672, GSM2180144, GSM2180145, GSM2180138, and GSM2180139) were run through the same pipeline to obtain expression levels over the same gene set. The other three datasets (GSM3408648, GSM3408667, and GSM3408685), only available as read counts per gene, were normalized to each gene’s total exonic length in order to be comparable to an RPKM-like expression density (to within an overall sample-dependent scaling factor). The data for each pair from among these 32 datasets were filtered to exclude snoRNA genes (some of which were expressed at high levels in a highly variable manner from sample to sample) and any genes that had precisely zero expression in either sample. This typically yielded ∼15,000–19,000 informative pairs of expressed genes per sample pair, for which Spearman correlations were calculated.

The BrainSpan dataset was obtained from http://www.brainspan.org/. These data altogether included 578 samples from 41 different donors spread across 30 ages ranging from 8 pcw through 40 years old and covering 26 brain regions. Expression levels (RPKM) in each BrainSpan sample were provided for 18,529 genes found in the RefSeq annotation for hg19 (GRCh37). The regions (a) ventrolateral prefrontal cortex (VFC), (b) medial prefrontal cortex (MFC), and (c) dorsolateral prefrontal cortex (DFC) included, respectively, 38, 37, and 39 BrainSpan samples at any age. The genome-wide expression profiles of the 7 SNR-derived organoid samples analyzed in **Supplementary Fig. 12** were analyzed with respect to hg38; for direct comparison to the BrainSpan samples in these regions, 19,889 hg38 genes were directly translatable to the hg19 RefSeq annotation, of which 18,448 genes overlapped genes documented by BrainSpan. A Spearman correlation coefficient was then calculated between each organoid sample and each of BrainSpan sample for each brain region, after removing genes with zero expression in either sample pair.

### Electrophysiology

#### Slice preparation

Acute slices for electrophysiology experiments were obtained from 5-month-old SNR-derived organoids. SNR-derived organoids were placed in ice-cold “cutting” artificial cerebrospinal fluid (ACSF), containing: KCl 2.5 mM, MgCl_2_ 7 mM, NaH_2_PO4 1.25 mM, choline-Cl 105 mM, NaHCO_3_ 25 mM, D-glucose 25 mM, Na^+^-pyruvate 3 mM, Na^+^-L-ascorbate 11 mM, and CaCl_2_ 0.5 mM. Tissue sections (200-µm-thick) were cut with a Leica VT1200 vibratome in ice-cold “cutting” ACSF. Sections were allowed to recover for 30 min in standard ACSF in a 37°C water bath before being stored at room temperature in standard ASCF. Standard ACSF contains: NaCl 124 mM, KCl 2.5 mM, MgCl_2_ 1 mM, NaH_2_PO_4_ 1.25 mM, NaHCO_3_ 25 mM, D-glucose 25 mM, Na^+^-pyruvate 1 mM, Na^+^-L-ascorbate 1 mM, and CaCl_2_ 2 mM. Both “cutting” and standard ACSF were bubbled with 95% O_2_/5% CO_2_.

#### Patch-clamp recordings

Individual slices were transferred to a recording chamber and submerged in standard ACSF bubbled with 95% O_2_/5% CO_2_. To characterize the electrophysiological properties of individual cells, whole-cell patch-clamp recording was performed at room temperature under constant perfusion of standard ACSF. Cells were visualized under a 40x water-immersion objective mounted on an upright microscope equipped with infrared differential interference contrast and a digital camera. Whole-cell patch-clamp recordings were performed using glass pipette electrodes (resistance 3–7 MΩ) pulled from borosilicate capillaries (BF150-86-10, Sutter Instruments, Novato, CA, USA) using a P-97 pipette puller (Sutter Instruments). Recording data were acquired using a Multiclamp 700B amplifier (Molecular Devices, Palo Alto, CA, USA) with Bessel filter of 2 kHz and digitized at 50 kHz using Axon DigiData 1500A (Molecular Devices). The Axon pClamp 10 (Molecular Devices) software was used to control the experiment. To characterize the intrinsic electrical properties of neurons, membrane voltage was recorded in the current clamp configuration and electrodes were filled with intracellular solution containing KMeS 123 mM, KCl 5 mM, HEPES 10 mM, Na-ascorbate 3 mM, MgCl_2_ 4 mM, Na_2_-phosphocreatine 10 mM, Na_2_GTP 0.4 mM, and Na_2_ATP 4 mM. 0.5% biocytin was added for a subset of experiments to visualize neuronal morphology. The resting membrane potential was recorded in the I=0 mode within 30 s after establishing the whole-cell patch. A small bias current was applied to maintain the baseline membrane voltage between -60 ∼ -75 mV. Square-pulse currents (1 or 2 s) were injected every 5 s, starting from hyperpolarizing amplitudes to increasingly more depolarized amplitudes, with a step size of 5 or 10 pA determined by the input resistance of the cell. For consistency, only the first 1 s of the current clamp data was analyzed. Measurements were not corrected for liquid junction potential. After patch-clamp recordings, sections were fixed in 4% paraformaldehyde (PFA) in PBS at 4°C overnight for subsequent immunohistochemical staining.

#### Current-clamp data analysis

To automate the analysis of current clamp data, we built a data analysis pipeline in Python using the DataJoint relational database (Yatsenko et al., 2015) and custom-written scripts adapted from the AllenSDK package (Allen SDK, 2015). The Axon binary files were converted into Python arrays using the stfio package from Stimfit (Guzman et al., 2014). We used the ephys module from AllenSDK as a baseline analysis framework and added methods to extract eletrophysiological features from our current clamp data. The analysis code was called within the DataJoint data processing pipeline to automatically analyze all recording data. The analysis results and experimental metadata were stored in the DataJoint relational database. Below we detail the analysis of each electrophysiological feature.

Rheobase sweep was determined as the first depolarizing sweep that has at least one action potential in the first 500 ms. Single action potential properties were measured using the first action potential of the rheobase sweep. Spike threshold was defined as the voltage where ΔV/Δt achieved 5% of the upstroke (maximum rising phase ΔV/Δt). Spike amplitude was measured as the difference between the peak voltage and the threshold voltage. Spike width was measured at the half-height of the spike. Spike trough (fast) was measured at 5 times spike width after the spike threshold. After-hyperpolarization (AHP) was defined as the difference between the trough voltage and the threshold voltage. First-spike latency was measured at 5 pA above the rheobase current. Since not all recordings have a current injection step size of 5 pA, the first-spike latency at rheobase + 5 pA was linearly interpolated using the two sweeps right above and below the target current.

The adaptation index was calculated on a subset of recordings that had at least 3 supra-threshold sweeps each with at least 4 spikes. Cells that did not meet these criteria had an undefined adaptation index. For each pair of consecutive inter-spike intervals (ISI), the adaption index was calculated as (ISI_2_ – ISI_1_) / (ISI_1_ + ISI_2_). Since different sweeps and recordings may have different numbers of spikes, we calculated the averaged adaptation index for each sweep using only the first two adaptation indices. We then averaged the adaptation indices of the first 3 supra-threshold sweeps as the adaptation index of each cell.

The F-I curve slope was calculated by a linear fit of firing rate against current injection amplitude. Data points after the maximum firing rate was reached were excluded from the fit. Cells with a maximum firing of only one spike per sweep were assigned a F-I slope of zero.

Input resistance was calculated using a linear fit of the maximum voltage deflection of each hyperpolarizing sweep against the current injection amplitude. The membrane time constant (tau) for each hyperpolarizing sweep was calculated by fitting a single exponential from 10%–100% of the maximum voltage deflection. Membrane capacitance was then calculated by first averaging the time constant of all hyperpolarizing sweeps and then dividing by input resistance.

The sag ratio for each hyperpolarizing sweep was calculated as (V_peak_ – V_steady-state_) / (V_peak_– V_baseline_), where V_peak_ is the maximum voltage deflection within the first 500 ms and V_steady-state_ is the averaged membrane voltage during the last 100 ms of the 1-s current injection. For each cell, two sag ratios with V_peak_ closest to -100 mV were averaged.

#### Hierarchical clustering

Using the above electrophysiological features, we constructed a cell-feature matrix. Hierarchical clustering was performed using the average linkage method and correlation distance metric. Briefly, each feature column was scaled to zero-mean and unit-variance, and the correlation distance was calculated on scaled features. For each pair of cells, if either of them had undefined adaptation index, the correlation was calculated without using adaptation index. Some features (spike width, first-spike latency, input resistance, and capacitance) had skewed distribution as judged by quantile-quantile plot and were log-transformed for distance calculation. After hierarchical clustering, the dendrogram tree was cut and 5 clusters was used for downstream analysis.

#### Synaptic current recordings

To measure synaptic currents, whole-cell voltage clamp was performed and electrodes were filled with intracellular solution containing CsMeS 120 mM, CsCl 5 mM, HEPES 10 mM, CaCl2 0.5 mM, EGTA 2 mM, Na-Ascorbate 3 mM, MgCl_2_ 4 mM, Glucose 5 mM, Na_2_-phosphocreatine10 mM, Na_2_GTP 0.4 mM, Na_2_ATP 4 mM. The membrane voltage was held at -70 mV for recording excitatory post-synaptic currents and +10 mV for recording inhibitory post-synaptic currents. The synaptic currents were analyzed using the Clampfit 10 software.

#### Western blotting

Protein lysates were prepared with Laemmli Buffer (Bio-Rad, 1610737) and loaded onto a sodium dodecyl sulfate (SDS)-polyacrylamide gel electrophoresis (PAGE) chamber to separate protein samples. Separated proteins were transferred to a polyvinylidene difluoride (PVDF) membrane overnight at 4°C. Following transfer, membranes were blocked with 5% milk in PBS with 0.1% Tween 20 (PBST) for 1–2 h while rocking at room temperature. Membranes were then incubated in primary anti-SHANK3 (Santa Cruz Biotechnology, sc-30193) at 1:350, or anti-TUJ1 (Covance, MMS-435P) at 1:38,000 in 5% milk in PBST and slowly rocked at 4°C overnight. Membranes were washed three times for 10 min with PBST and then incubated with horseradish peroxidase (HRP)-conjugated secondary antibody (HRP anti-rabbit [Jackson ImmunoResearch, 111-035-144] or HRP anti-mouse [Vector Laboratories, PI-2000) at 1:15,000-1:25,000) in 5% milk in PBST for 1–2 h at room temperature. After three 10-min washes in PBST, the membrane was placed in deionized water and gently dried before adding chemiluminescent HRP substrate (EMD Millipore, WBKLS0100) for visualizing proteins. Blotting X-ray film (Genesee Scientific, 30-507L) was then placed over the membrane with a sheet protector in between. The film was processed using a Konica SRX-101 machine. Films were imaged with ChemiDoc XRS+ System with Image Lab Software (Bio-Rad, 1708265) and analyzed using ImageJ.

### Immunohistochemistry and imaging

#### SNR-derived organoids

Rosettes were fixed in 4% paraformaldehyde for 15 min at room temperature, washed three times with PBS, followed by permeabilization with 0.3% Triton-X for 15 minutes at room temperature. Rosettes were then blocked in 3% BSA with or without 10% Normal Goat Serum for 1 hour at room temperature. Primary antibodies (**Supplementary Table 7**) were diluted in 3% BSA and applied over night at 4°C. Following primary antibody incubation, rosettes were washed three times with PBS. Secondary antibodies were diluted in 3% BSA and applied for 2 hours at room temperature in the dark. Rosettes were then washed three times with PBS and incubated with Hoechst 33342 nuclear counterstain (1:50,000 in PBS) for 3 min at room temperature.

SNR-derived organoids were fixed in 4% paraformaldehyde overnight at 4°C and washed three times with PBS before passing through a sequential sucrose gradient of 10%, 20%, and 30% sucrose solutions. SNR-derived organoids were then placed in cryosectioning molds, embedded in OCT and flash frozen, then stored at -80°C. SNR-derived organoids were cryosectioned using a Leica Cryostat machine, and sections were cut at 10-20 µm thickness and adhered to positively-charged microscope slides. Adhered cryosections were then processed for immunostaining as described above. All samples were mounted with coverslips using Aqua Poly Mounting Media (Polysciences, Inc.). Fluorescent imaging was performed at the University of Utah Cell Imaging Core Facility using Nikon A1, Nikon A1R, or Zeiss 880 Airyscan confocal microscopes. Data processing and quantification was performed using Fiji, Volocity 5.1, and/or CellProfiler by setting the threshold to remove background and identify cell bodies or nuclei as objects. Cell percentages were obtained by counting the number of objects staining positive for a marker as a ratio of the total number of nuclear objects identified by Hoechst staining. For quantification of MAP2-expressing cells in control and SHANK3-deficient organoids, images were analyzed by two independent investigators blinded to the genotypes. For quantification of synaptic puncta, *Shank3*-/- adult mouse brain tissue was used as negative control to optimize the threshold for detecting SHANK3.

#### Primary human fetal brain tissue

Postmortem human brain specimens were obtained from University of Cambridge, UK upon pregnancy terminations. All procedures were approved by the research ethical committees and research services division of the University of Cambridge and Addenbrooke’s Hospital in Cambridge (protocol 96/85, approved by Health Research Authority, Committee East of England—Cambridge Central in 1996 and with subsequent amendments, with the latest approved November 2017). Tissue was handled in accordance with ethical guidelines and regulations for the research use of human brain tissue set forth by the National Institute of Health (NIH) (http://bioethics.od.nih.gov/humantissue.html) and the World Medical Association Declaration of Helsinki (http://www.wma.net/en/30publications/10policies/b3/index.html). Embryonic and fetal age was extrapolated based on the last menstrual period of the mother and crown to rump length (CRL) measured by the clinicians. The tissues were dissected, fixed (24 h, 4°C) in 4% paraformaldehyde (PFA) and incubated in 30% sucrose for 24 h at 4°C. Brains were than snap frozen in optimal cutting temperature (OCT) medium, sectioned (15 μm) using a cryostat (Leica, Germany) and stored at − 80 °C. Sections were thawed just prior to staining and rinsed in PBS. Fixed tissues were exposed to antigen retrieval step with 1 mM EDTA solution for 60 min at 60°C. After washing using 1X PBS, tissues were blocked and permeabilized using 5% normal goat serum (NGS; Vector) plus 0.1% Triton X-100 in PBS for 1 h at room temperature. After washing with PBS plus 0.1% Tween-20, cells were incubated overnight at 4 °C with primary antibodies (GSX2: rabbit 1:200, Merck-Millipore ABN162; PAX6 mouse 1:80, DSHB) diluted in solution containing 2.5% NGS and 0.02% Triton X-100. Alexa Fluor secondary antibodies (ThermoFisher scientific) were used 1:1000 in PBS for 1 h RT, followed by 3X washes with PBST and 10 min incubation at RT with 1:10000 of DAPI (ThermoFisher scientific) in PBS. Images were acquired with Leica confocal TCS SP5 (Leica Microsystem). For RNA-FISH, 9 weeks human fetal brain sections were dehydrated in 5 min steps in 70% and 100% ethanol, respectively. A hydrophobic barrier was created around the sections and air dried for 10 min. Sections were incubated with RNAscope® Hydrogen Peroxide solution (ACD, 322335) for 10 min at RT. After incubation, slides were washed twice with ddH_2_O. Protease plus (ACD, 322331) was added to the samples and incubated in a HybEZ oven for 30 minutes at 40°C. After incubation, slides were washed in ddH_2_O and RNAscope in situ hybridizations were performed according to the manufacturer’s instructions, using the RNAscope® Multiplex Fluorescent Assay v2 (Advanced Cell Diagnostics) for fixed frozen tissue. Following probes were used (indicated with gene target name for human, respective channel, and catalogue number all Advanced Cell Diagnostics): Pax6 (Ch1, 588881) and Gsx2 (Ch4, 540601). Probe hybridization took place for 2 h at 40 °C and slides were then rinsed in 1 × wash buffer, followed by four amplification steps (according to protocol). Brain sections were then labelled with DAPI, and mounted with Prolong Gold mounting medium (P36930, Thermo Fisher Scientific). Slides were stored at 4°C before image acquisition using a Nikon Eclipse Ti-E microscope (Nikon Instruments).

## ACKNOWLEDGEMENT

The authors are thankful to Dr. Ricardo Dolmetsch and Dr. Carl Ernst for sharing control iPSC lines; to Dr. Theo Palmer for assistance with material transfer; to Laura Bell for help with immunostaining, imaging, and image quantifications; to Dr. Jianmin Zhang for help with cryosectioning; to Oleksandr Stepanenko, Dmitro Zhmurko, Roman Savitskyi, David Richardson, Trevor Tanner, and Dr. Thomas Cheatham for help with web-browser; to Dr. Jian Zhou for discussion of single-cell RNA-seq analysis methods; to Dr. Brian Dalley, Opal Allen, Chris Stubben, Brian Lohman and Utah high-throughput genomics facility for help with sequencing and data analysis; to Utah cell imaging facility for help with imaging; to Dr. Megan Williams, Dr. Monica Vetter, Dr. Sungjin Park, and Dr. Richard Dorsky for sharing reagents and commenting on the manuscript; to Dr. Yong-Hui Jiang and Dr. Xiaoming Wang for sharing WT and *SHANK3*-/- mouse brain tissue. This work was supported by grants from the NIMH, NINDS, Brain Research Foundation, Brain and Behaviour Research Foundation, Whitehall Foundation, and Utah Neuroscience and Genome Project Initiatives.

