## Supplementary Figures for "Modeling autism-associated SHANK3 deficiency using human cortico-striatal organoids generated from single neural rosettes"

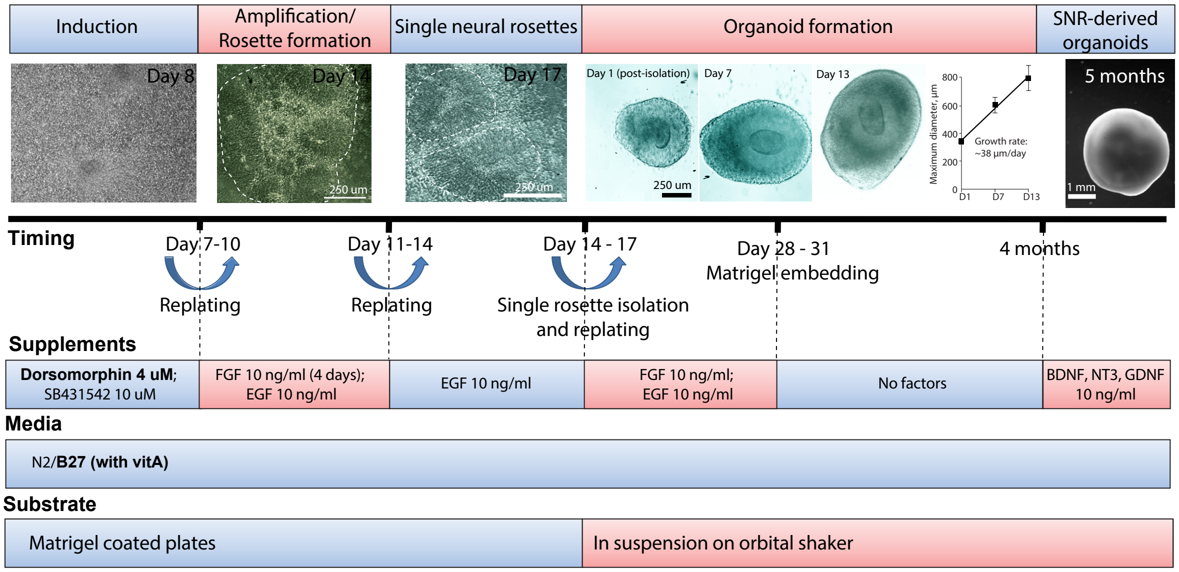

**Supplementary Figure 1. Organoid generation protocol.** Dual SMAD inhibitors (Dorsomorphin and SB431542) are used to convert pluripotent stem cells into neuroepithelial-like cells (Induction). FGF and EGF are used to amplify the population of neural progenitors and to induce the formation of neural rosettes (Amplification/Rosette formation). SNRs (200-250 μm in diameter) are isolated 14-17 days post induction (Single neural rosettes) and cultured in 1:1 mixture of N2 and B27 (with vitamin A) media supplemented with EGF and FGF on an orbital shaker in an incubator (5% $CO_2$ and 37°C) (Organoid formation). 28-31 days post-induction organoids are embedded in Matrigel and cultured on orbital shaker in N2/B27 (with vitamin A) media without growth or trophic factors. 4 months post-induction, trophic factors are added to the media to promote functional and synaptic neuronal maturation (SNR-derived organoids). Additional protocol details are provided in the materials and methods section. Images are also presented in Fig. 1 and 3.

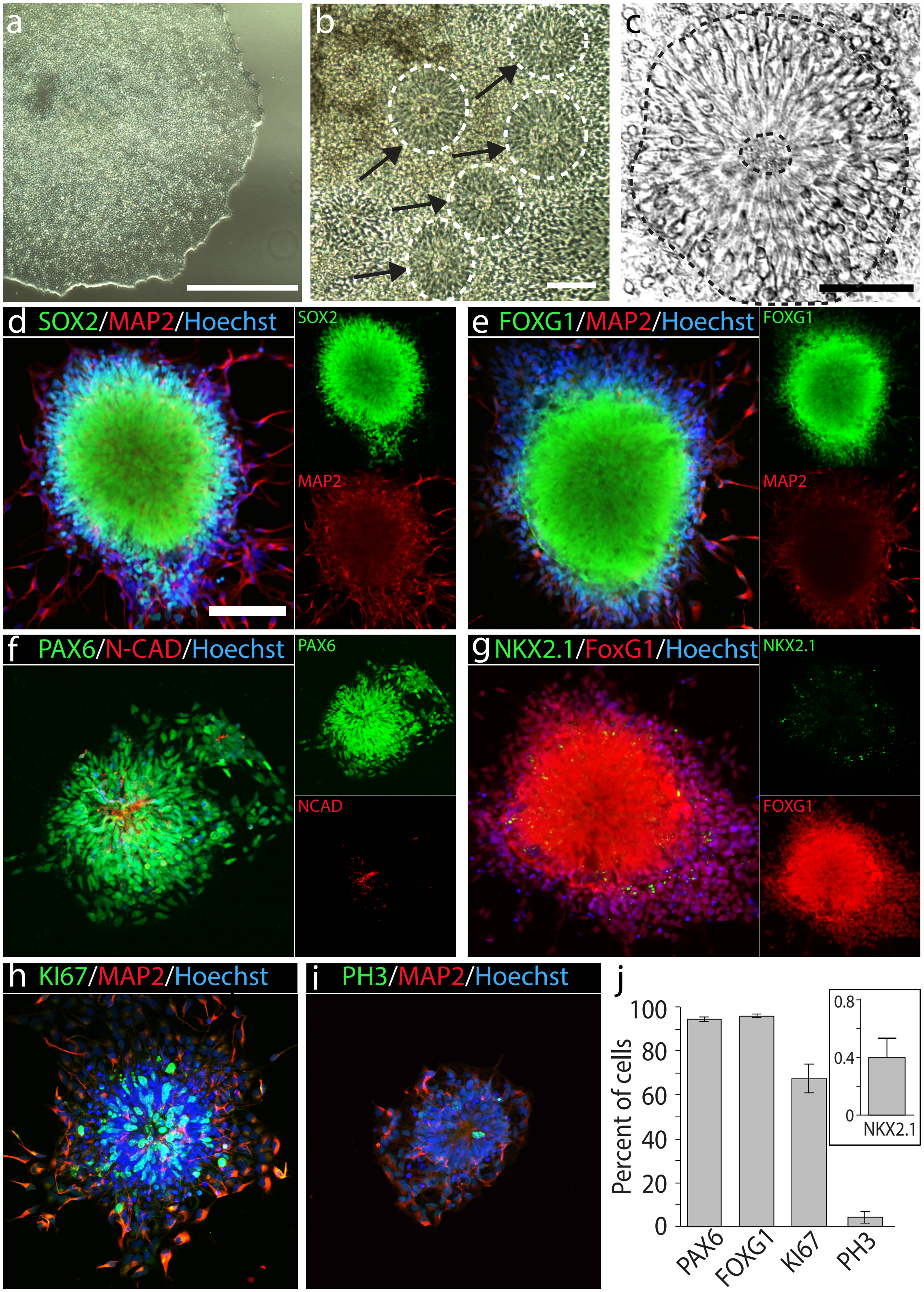
a
b
c
d SOX2/MAP2/Hoechst
SOX2
MAP2
e FOXG1/MAP2/Hoechst
FOXG1
MAP2
f PAX6/N-CAD/Hoechst
PAX6
NCAD
g NKX2.1/FoxG1/Hoechst
NKX2.1
FOXG1
h KI67/MAP2/Hoechst
i PH3/MAP2/Hoechst
j
Percent of cells
100
80
60
40
20
0
PAX6
FOXG1
KI67
PH3
0.8
0.4
0
NKX2.1

**Supplementary Figure 2. Generation and characterization of stem cell-derived SNRs.**

**a-b**, Low magnification images of an iPSC colony (**a**) and cluster of iPSC-derived SNRs (**b**). **c**, High magnification image of an isolated SNR. **d-h**, Images of isolated SNRs immunostained with antibodies against SOX2 and MAP2 (**d**), FOXG1 and MAP2 (**e**), PAX6 and N-Cad (**f**), PAX6 and N-Cad (**g**), Ki67 and MAP2 (**h**), and PH3 and MAP2 (**i**). **j**, Percent of cells in SNRs expressing PAX6 (n = 4 rosettes), FOXG1 (n = 4), Ki67 (n = 3), and PH3 (n = 4), and NKX2.1 (n = 8). Data presented as means ± s.e.m. Scale bars = 1000 (**a**), 200 (**b**), 50 (**c**), and 100 μm (**d**).

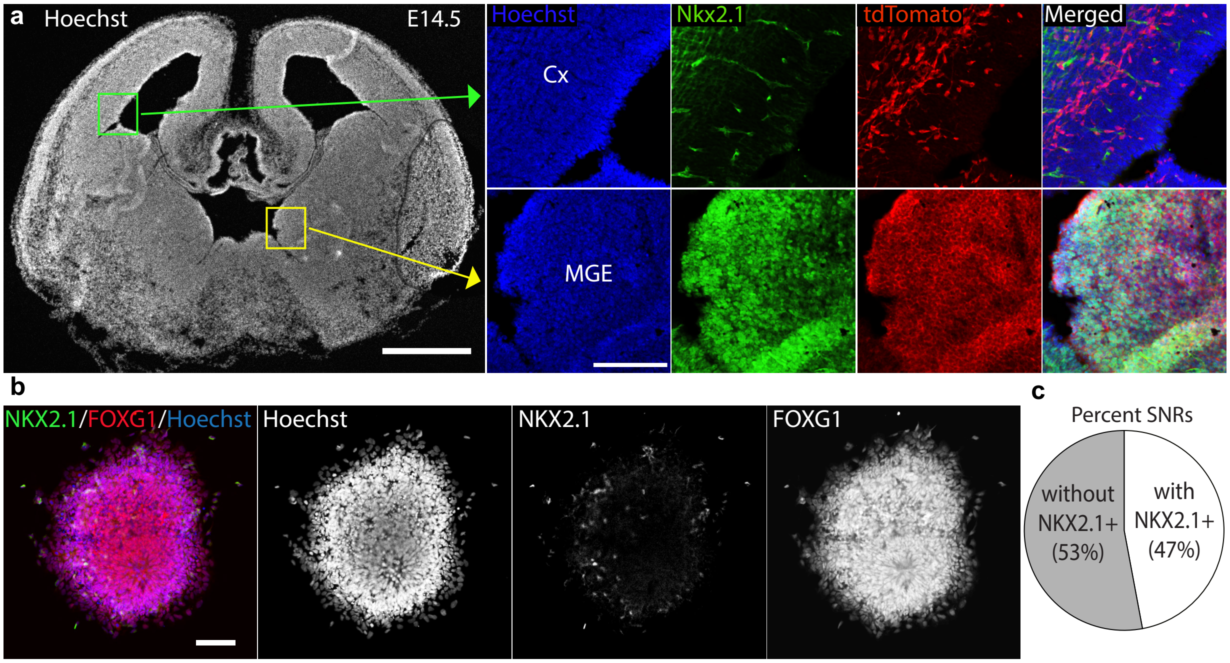

a
Hoechst
E14.5
Hoechst
Nkx2.1
tdTomato
Merged
Cx
MGE
b
NKX2.1/FOXG1/Hoechst
Hoechst
NKX2.1
FOXG1
c
Percent SNRs
without NKX2.1+ (53%)
with NKX2.1+ (47%)

**Supplementary Figure 3. Characterization of NKX2.1 expression in SNRs as compared to developing mouse brain.** **a**, Image of a coronal brain section from a conditional $Rosa^{tdTomato}$/$Nkx2.1^{Cre}$ mouse at E14.5 (Xie et al., Plos Biol., 2017) immunostained with antibodies against Nkx2.1 and DSRed to validate anti-Nkx2.1 antibody specificity. Nkx2.1 expression was detected in the ventral, but not dorsal, progenitors. **b**, Image of an isolated SNR immunostained with antibodies against NKX2.1 and FOXG1. **c**, Percent of SNRs with and without NKX2.1-expressing cells (n = 17 rosettes). Scale bars = 500 (**a**), and 100 µm (**a** [zoom in] and **b**).

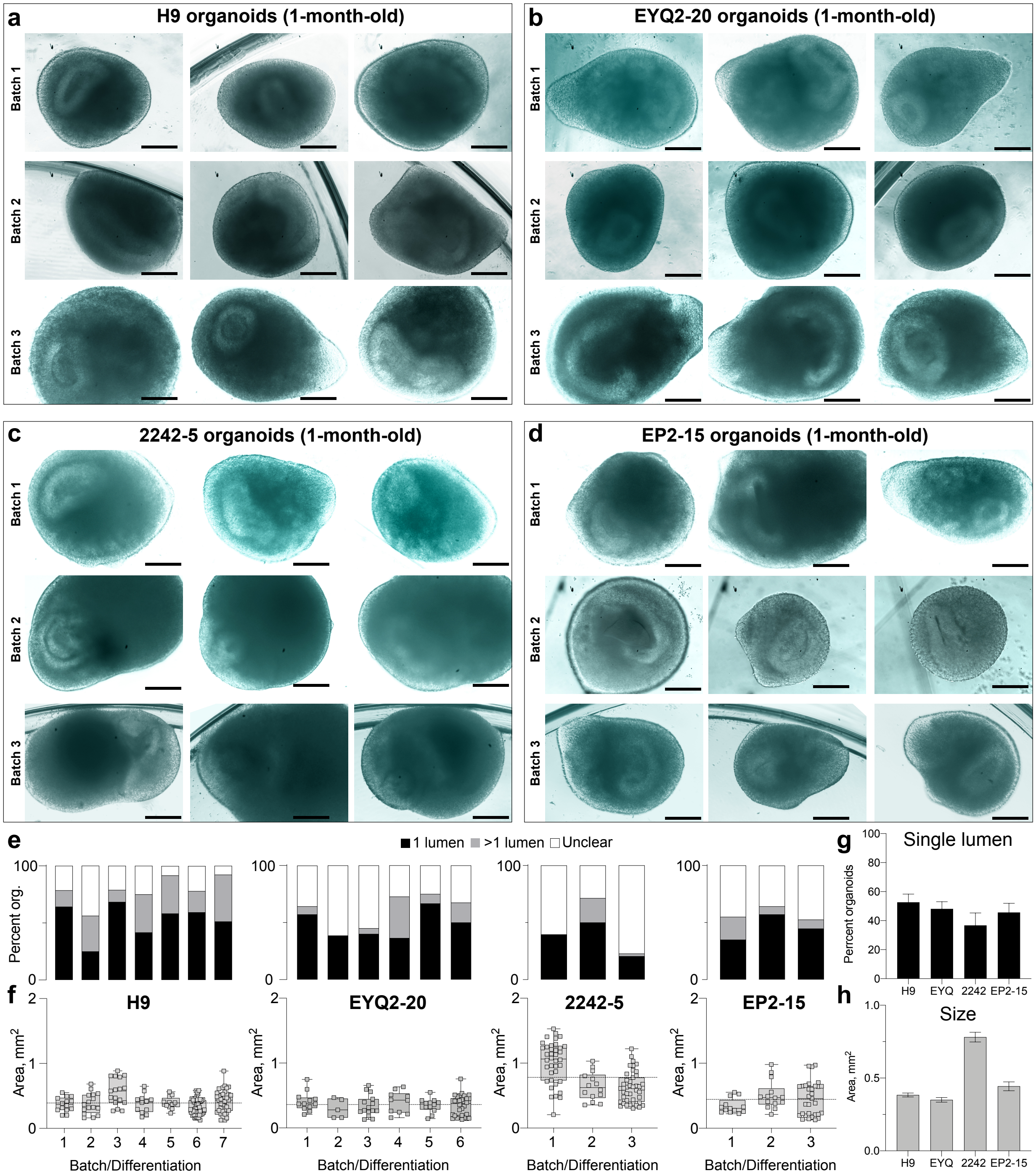

a
H9 organoids (1-month-old)
b
EYQ2-20 organoids (1-month-old)
c
2242-5 organoids (1-month-old)
d
EP2-15 organoids (1-month-old)
Batch 1
Batch 2
Batch 3
e
1 lumen
>1 lumen
Unclear
Percent org.
f
H9
EYQ2-20
2242-5
EP2-15
Area, mm$^2$
Batch/Differentiation
g
Single lumen
Percent organoids
h
Size
H9
EYQ
2242
EP2-15

**Supplementary Figure 4. Characterization of growth and organization of 1-month-old SNR-derived organoids. a-d**, Representative images of SNR-derived organoids produced from four pluripotent stem cell lines (H9, EYQ2-20, 2242-5, and EP2-15) in different differentiation batches. **e-h,** Quantification of lumen (**e, g**) and size (**f, h**) of organoids by differentiation batch. Data presented as individual data point, median, and min and max values (whiskers). **f**, Quantification of lumen and size of organoids by line (H9: n=159 organoids/7 batches; EYQ2-20: n=84 organoids/6 batches; 2242-5: n=95 organoids/3 batches; EP2-15: n=58 organoids/3 batches). Data presented as means ± s.e.m. Scale bars = 250 μm.

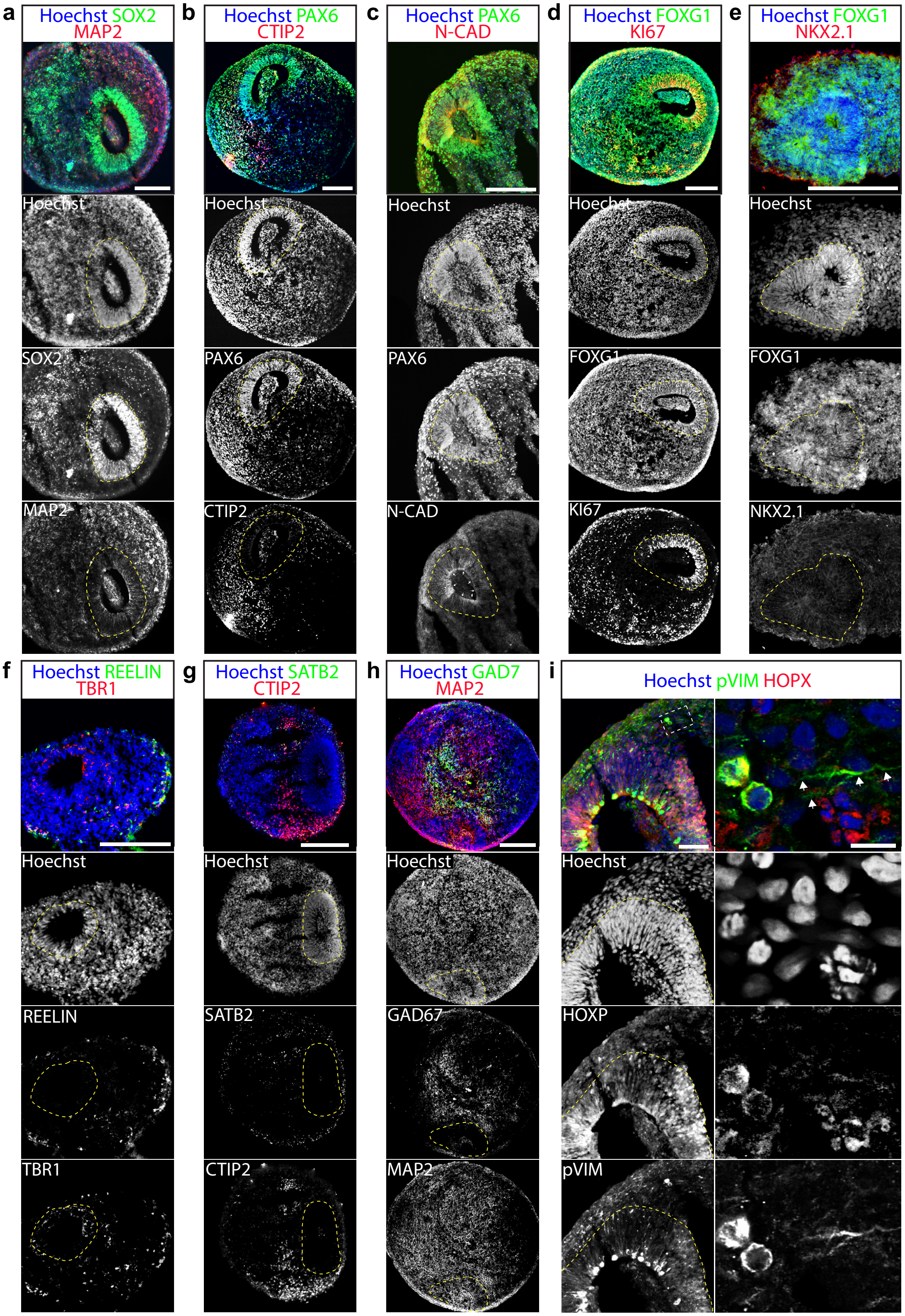
a
Hoechst SOX2 MAP2
Hoechst
SOX2
MAP2
b
Hoechst PAX6 CTIP2
Hoechst
PAX6
CTIP2
c
Hoechst PAX6 N-CAD
Hoechst
PAX6
N-CAD
d
Hoechst FOXG1 KI67
Hoechst
FOXG1
KI67
e
Hoechst FOXG1 NKX2.1
Hoechst
FOXG1
NKX2.1
f
Hoechst REELIN TBR1
Hoechst
REELIN
TBR1
g
Hoechst SATB2 CTIP2
Hoechst
SATB2
CTIP2
h
Hoechst GAD7 MAP2
Hoechst
GAD67
MAP2
i
Hoechst pVIM HOPX
Hoechst
HOXP
pVIM

**Supplementary Figure 5. Representative images of organoid sections immunostained with antibodies against different cell-type specific markers.** **a**, SOX2 and MAP2. **b**, PAX6 and CTIP2. **c**, PAX6 and N-Cad. **d**, FOXG1 and Ki67. **e**, FOXG1 and NKX2.1. **f**, REELIN and TBR1. **g**, SATB2 and CTIP2. **h**, GAD67 and MAP2. **i**, pVIM and HOPX. Scale bars = 200 μm (**a-h**) and 50 and 10 μm (**i**).

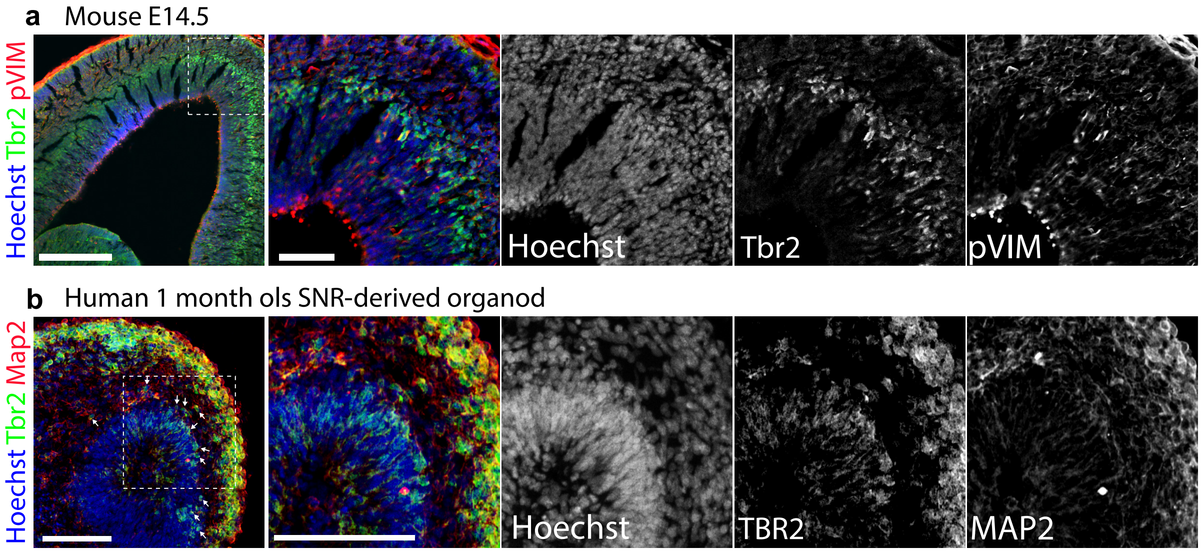
a
Mouse E14.5
Hoechst Tbr2 pVIM
Hoechst
Tbr2
pVIM
b
Human 1 month ols SNR-derived organod
Hoechst Tbr2 Map2
Hoechst
TBR2
MAP2

**Supplementary Figure 6. Representative images of organoid sections immunostained with antibodies against TBR2 in comparison to embryonic mouse cortex.** **a**, Image of a coronal embryonic mouse brain (E14.5) section immunostained with antibodies against Tbr2 and pVIM. Tbr2 expression was detected in the subventricular zone. **b**, Image of an organoid section immunostained with antibodies against TBR2 and MAP2. Scale bars = 100 and 50 µm.

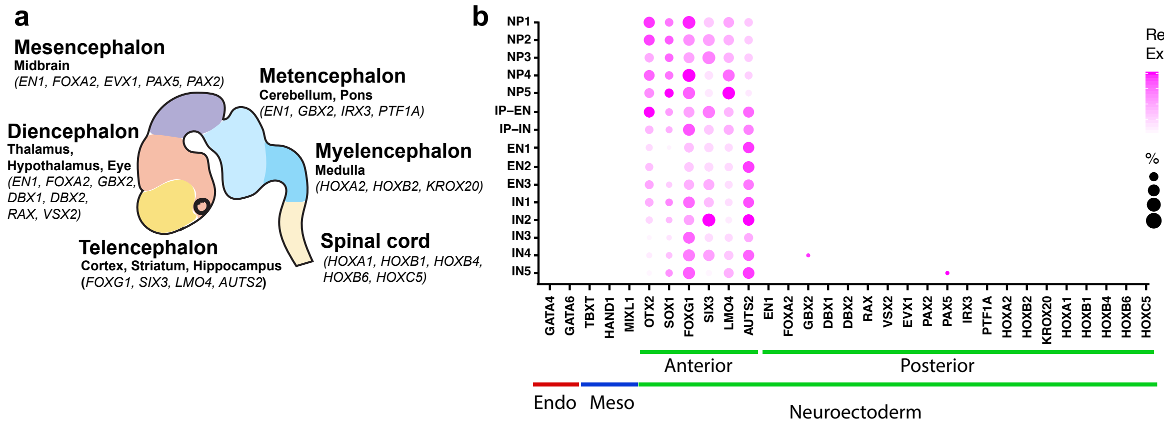
a
Mesencephalon
Midbrain
(EN1, FOXA2, EVX1, PAX5, PAX2)
Metencephalon
Cerebellum, Pons
(EN1, GBX2, IRX3, PTF1A)
Diencephalon
Thalamus,
Hypothalamus, Eye
(EN1, FOXA2, GBX2,
DBX1, DBX2,
RAX, VSX2)
Myelencephalon
Medulla
(HOXA2, HOXB2, KROX20)
Telencephalon
Cortex, Striatum, Hippocampus
(FOXG1, SIX3, LMO4, AUTS2)
Spinal cord
(HOXA1, HOXB1, HOXB4,
HOXB6, HOXC5)
b
NP1
NP2
NP3
NP4
NP5
IP–EN
IP–IN
EN1
EN2
EN3
IN1
IN2
IN3
IN4
IN5
GATA4
GATA6
TBXT
HAND1
MIXL1
OTX2
SOX1
FOXG1
SIX3
LMO4
AUTS2
EN1
FOXA2
GBX2
DBX1
DBX2
RAX
VSX2
EVX1
PAX2
PAX5
IRX3
PTF1A
HOXA2
HOXB2
KROX20
HOXA1
HOXB1
HOXB4
HOXB6
HOXC5
Anterior
Posterior
Endo
Meso
Neuroectoderm
Relative
Express.
1.00
0.75
0.50
0.25
0.00
% cells
25
50
75
100

**Supplementary Figure 7. Expression of endodermal, mesodermal, and neuroectodermal markers in 1-month-old SNR-derived organoids. a,** Cartoon visualization of the embryonic mammalian brain with brain region-specific markers. **b,** Dotplot visualization of region- and cell-type specific marker expression in different cell clusters.

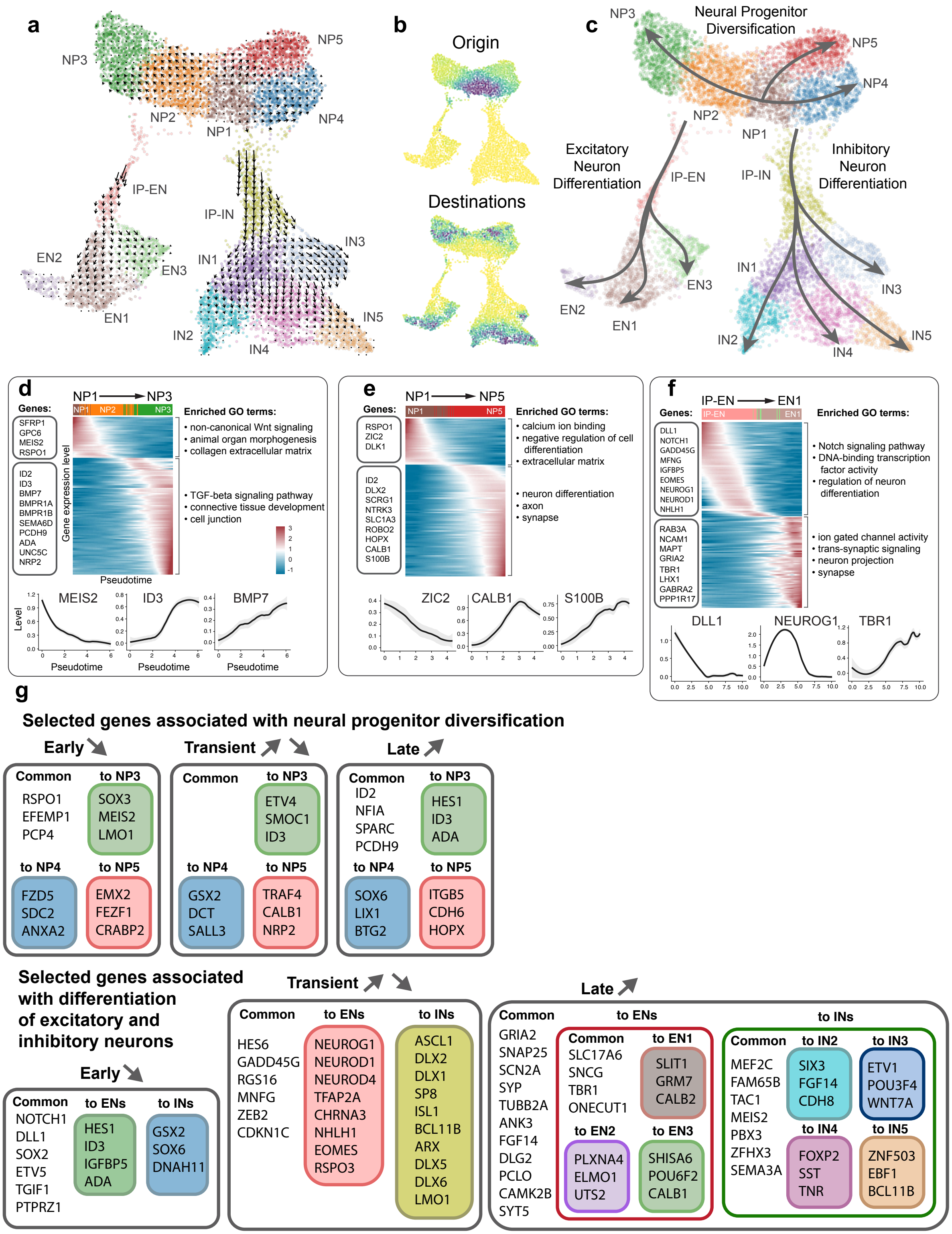

a
NP3
NP5
NP2
NP1
NP4
IP-EN
IP-IN
EN2
EN3
EN1
IN1
IN3
IN2
IN4
IN5
b
Origin
Destinations
c
NP3
Neural Progenitor Diversification
NP5
NP4
NP2
NP1
Excitatory Neuron Differentiation
IP-EN
IP-IN
Inhibitory Neuron Differentiation
IN1
IN3
EN3
EN2
EN1
IN2
IN4
IN5
d
NP1 → NP3
Genes:
SFRP1 GPC6 MEIS2 RSPO1
ID2 ID3 BMP7 BMPR1A BMPR1B SEMA6D PCDH9 ADA UNC5C NRP2
Gene expression level
Pseudotime
Enriched GO terms:
• non-canonical Wnt signaling
• animal organ morphogenesis
• collagen extracellular matrix
• TGF-beta signaling pathway
• connective tissue development
• cell junction
MEIS2
ID3
BMP7
Level
Pseudotime
e
NP1 → NP5
Genes:
RSPO1 ZIC2 DLK1
ID2 DLX2 SCRG1 NTRK3 SLC1A3 ROBO2 HOPX CALB1 S100B
Enriched GO terms:
• calcium ion binding
• negative regulation of cell differentiation
• extracellular matrix
• neuron differentiation
• axon
• synapse
ZIC2
CALB1
S100B
f
IP-EN → EN1
Genes:
DLL1 NOTCH1 GADD45G MFNG IGFBP5 EOMES NEUROG1 NEUROD1 NHLH1
RAB3A NCAM1 MAPT GRIA2 TBR1 LHX1 GABRA2 PPP1R17
Enriched GO terms:
• Notch signaling pathway
• DNA-binding transcription factor activity
• regulation of neuron differentiation
• ion gated channel activity
• trans-synaptic signaling
• neuron projection
• synapse
DLL1
NEUROG1
TBR1
g
Selected genes associated with neural progenitor diversification
Early
Common: RSPO1 EFEMP1 PCP4
to NP3: SOX3 MEIS2 LMO1
to NP4: FZD5 SDC2 ANXA2
to NP5: EMX2 FEZF1 CRABP2
Transient
Common
to NP3: ETV4 SMOC1 ID3
to NP4: GSX2 DCT SALL3
to NP5: TRAF4 CALB1 NRP2
Late
Common: ID2 NFIA SPARC PCDH9
to NP3: HES1 ID3 ADA
to NP4: SOX6 LIX1 BTG2
to NP5: ITGB5 CDH6 HOPX
Selected genes associated with differentiation of excitatory and inhibitory neurons
Early
Common: NOTCH1 DLL1 SOX2 ETV5 TGIF1 PTPRZ1
to ENs: HES1 ID3 IGFBP5 ADA
to INs: GSX2 SOX6 DNAH11
Transient
Common: HES6 GADD45G RGS16 MNFG ZEB2 CDKN1C
to ENs: NEUROG1 NEUROD1 NEUROD4 TFAP2A CHRNA3 NHLH1 EOMES RSPO3
to INs: ASCL1 DLX2 DLX1 SP8 ISL1 BCL11B ARX DLX5 DLX6 LMO1
Late
Common: GRIA2 SNAP25 SCN2A SYP TUBB2A ANK3 FGF14 DLG2 PCLO CAMK2B SYT5
to ENs
Common: SLC17A6 SNCG TBR1 ONECUT1
to EN1: SLIT1 GRM7 CALB2
to EN2: PLXNA4 ELMO1 UTS2
to EN3: SHISA6 POU6F2 CALB1
to INs
Common: MEF2C FAM65B TAC1 MEIS2 PBX3 ZFHX3 SEMA3A
to IN2: SIX3 FGF14 CDH8
to IN3: ETV1 POU3F4 WNT7A
to IN4: FOXP2 SST TNR
to IN5: ZNF503 EBF1 BCL11B

**Supplementary Figure 8. Developmental trajectories of 1-month-old SNR-derived organoids. a**, RNA velocity vector fields. Arrows indicate the extrapolated future states of cells in the UMAP plot. **b**, The origin and end states of the differentiation landscape determined using Markov random walk simulation on the velocity field. The density of cells at the end of the simulation is shown using the color scale ranging from yellow (low) to blue (high). **c**, Branching lineage trajectories constructed using Slingshot and initialized using the origin and end states of the Markov simulation. **d-f**, Smoothed gene expression heat maps of the top 150 genes differentially expressed along NP1->NP3 (**d**), NP1->NP5 (**e**), and IP-EN->EN1 (**f**) developmental trajectories. The genes are ordered by peak expression time on the pseudotime axis. Selected example genes are listed on the left and shown at the bottom. The top gene ontology terms are listed on the right. **g**, Examples of common and lineage-specific genes that showed differential expression along different differentiation trajectories (top: diversification of NPs, bottom: neuronal differentiation). The selected genes are divided into three categories: early, transient, and late, based on the timing for peak expression.

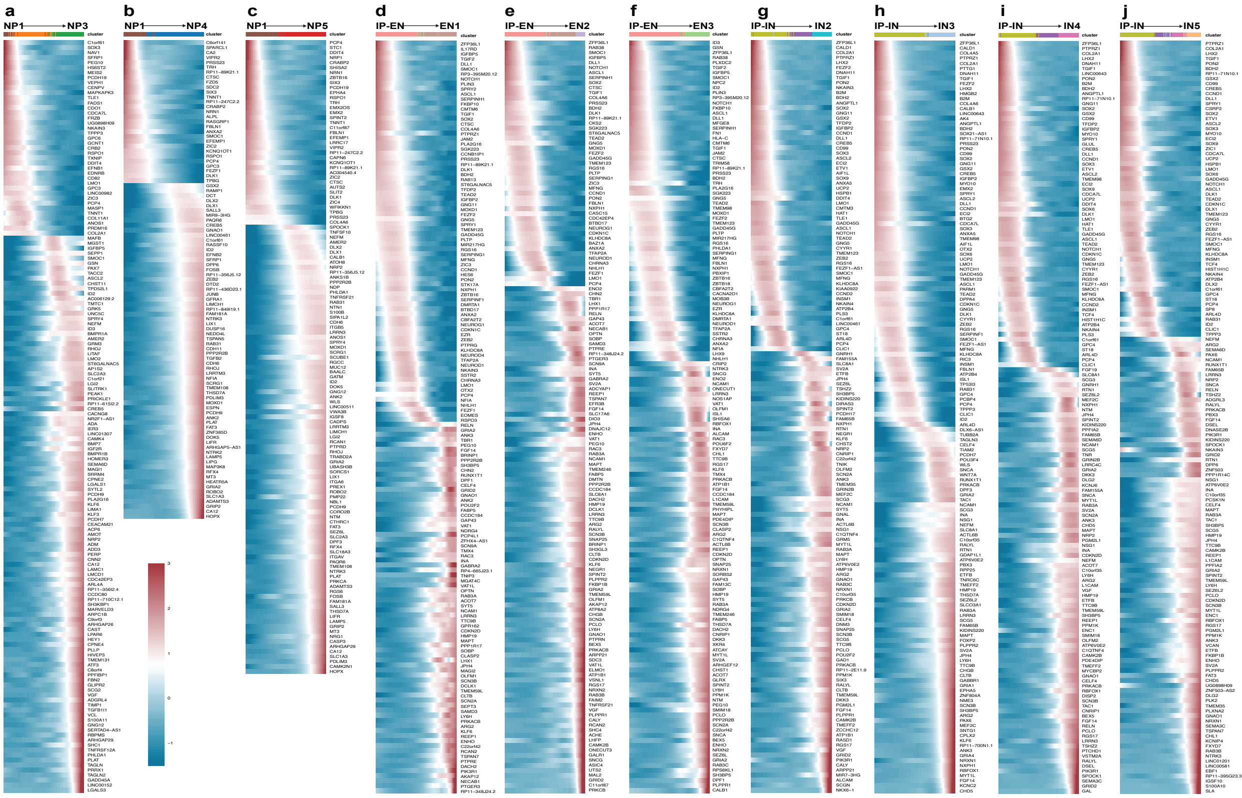

a
NP1 → NP3
b
NP1 → NP4
c
NP1 → NP5
d
IP-EN → EN1
e
IP-EN → EN2
f
IP-EN → EN3
g
IP-IN → IN2
h
IP-IN → IN3
i
IP-IN → IN4
j
IP-IN → IN5

**Supplementary Figure 9. Trajectory-specific genes. a**, NP1 -> NP3; **b**, NP1 -> NP4; **c**, NP1 -> NP5; **d**, IP-EN -> EN1; **e**, IP-EN -> EN2; **f**, IP-EN -> EN3; **g**, IP-IN -> IN2; **h**, IP-IN -> IN3; **i**, IP-IN -> IN4; and **j**, IP-IN -> IN5 trajectories. Genes showing differential expression displayed vertically. Horizontal axis displays pseudotime.

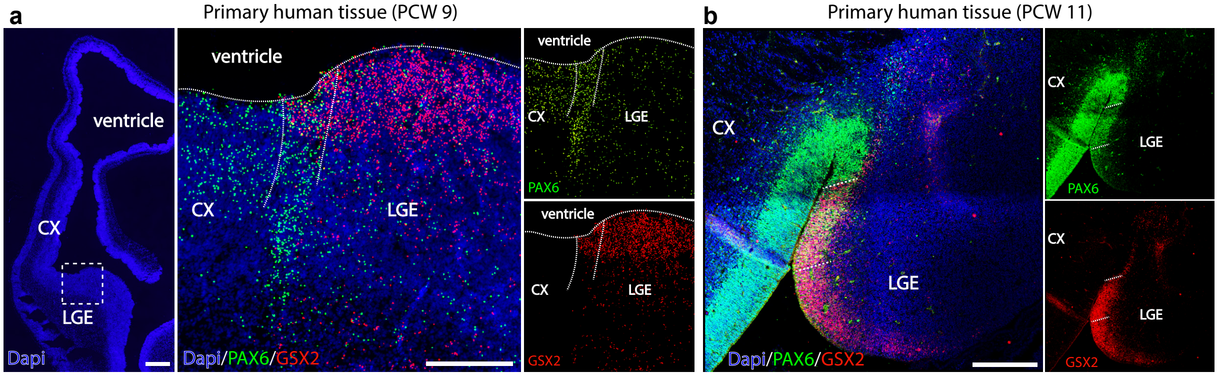
a
Primary human tissue (PCW 9)
ventricle
CX
LGE
Dapi
ventricle
CX
LGE
Dapi/PAX6/GSX2
ventricle
CX
LGE
PAX6
ventricle
CX
LGE
GSX2
b
Primary human tissue (PCW 11)
CX
LGE
Dapi/PAX6/GSX2
CX
LGE
PAX6
CX
LGE
GSX2

**Supplementary Figure 10. PAX6 and GSX2 expression in primary human tissue. a-b**, Images of human embryonic brain tissue sections, containing both pallial (CX) and subpallial regions (LGE), stained with FISH probes against *PAX6* and *GSX2* (**a**) or immunostained with antibodies against PAX6 and GSX2 (**b**). Scale bars = 500 and 200 µm (**a**) and 200 µm (**b**).

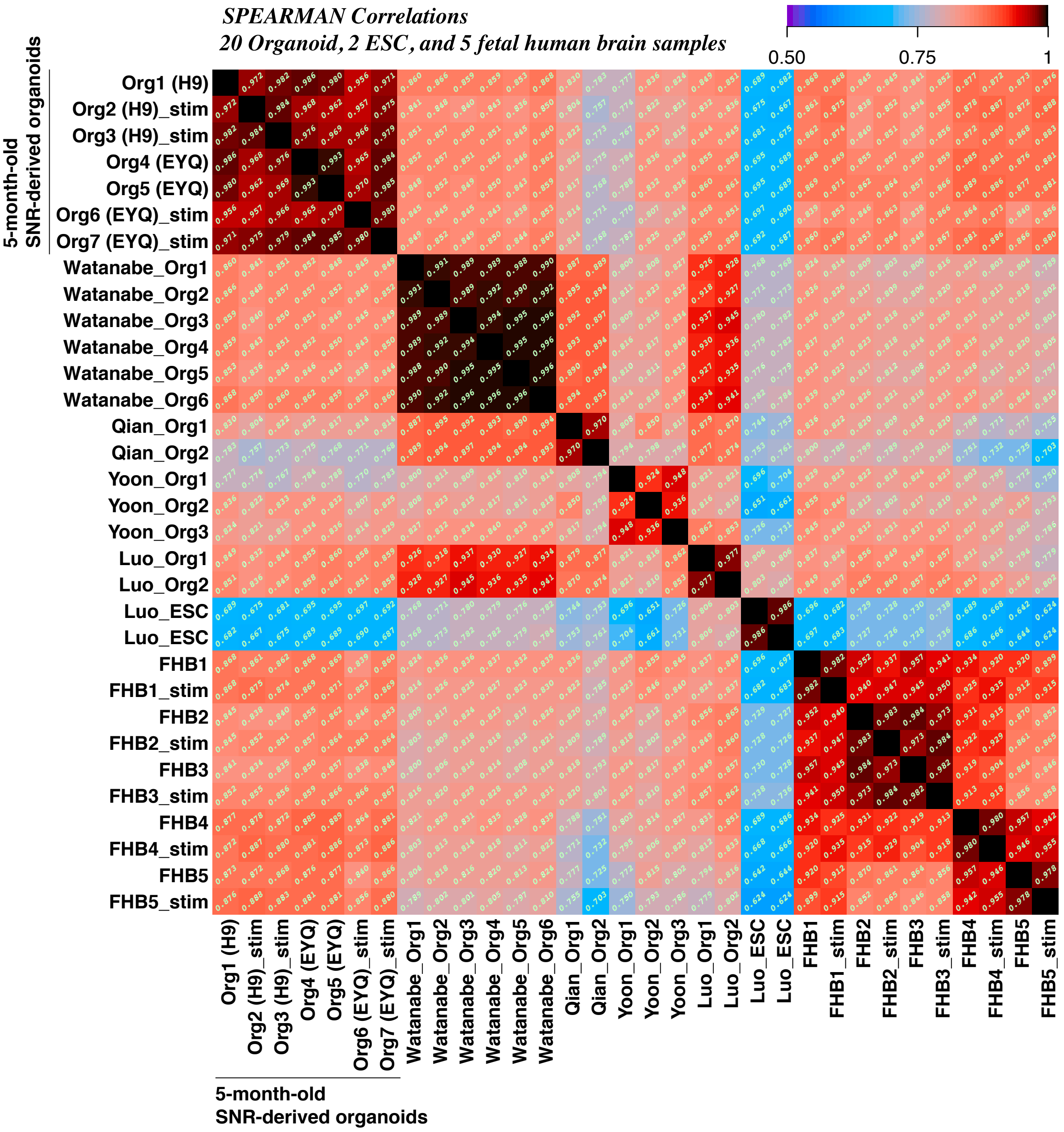
SPEARMAN Correlations
20 Organoid, 2 ESC, and 5 fetal human brain samples
0.50
0.75
1
5-month-old SNR-derived organoids
Org1 (H9)
Org2 (H9)_stim
Org3 (H9)_stim
Org4 (EYQ)
Org5 (EYQ)
Org6 (EYQ)_stim
Org7 (EYQ)_stim
Watanabe_Org1
Watanabe_Org2
Watanabe_Org3
Watanabe_Org4
Watanabe_Org5
Watanabe_Org6
Qian_Org1
Qian_Org2
Yoon_Org1
Yoon_Org2
Yoon_Org3
Luo_Org1
Luo_Org2
Luo_ESC
Luo_ESC
FHB1
FHB1_stim
FHB2
FHB2_stim
FHB3
FHB3_stim
FHB4
FHB4_stim
FHB5
FHB5_stim
5-month-old SNR-derived organoids

**Supplementary Figure 11. Transcriptional analysis to assess reproducibility.** Spearman correlations of expression profiles among 20 organoids ("Org"), 2 embryonic stem cell samples ("ESC"), and 10 fetal human brain ("FHB") samples. Pairwise correlation values shown by color scale and displayed in each off-diagonal cell. The following samples were used for analysis:

5-month-old SNR-derived telencephalic organoids: 7 biological replicates produced from H9 and EYQ stem cell lines, unstimulated or stimulated ("stim") with KCl for 6 hours;

2-month-old cortical organoids (Watanabe et al., Cell Reports 2017): 6 biological replicates produced from H9 stem cell line (GEO accession numbers: GSM2580319 [Org1], GSM2580321 [Org2], GSM2580323 [Org3], GSM2580325 [Org4], GSM2580327 [Org5], GSM2580329 [Org6]);

3.5-month-old telencephalic organoids (Qian et al., Cell 2016): 2 biological replicates produced from an iPSC stem cell line (GEO accession numbers: GSM2112671 [Org1] and GSM2112672 [Org2]);

3.5-month-old cortical spheroids (Joon et al., Nature Methods 2019): 3 biological replicates produced from three different iPSC stem cell lines (GEO accession numbers: GSM3408648 [Org1], GSM3408667 [Org2], and GSM3408685 [Org3]);

2-month-old cerebral organoids (Luo et al., Cell Reports 2016): 2 biological replicates produced from H9 stem cell line (GEO accession numbers: GSM2180144 [Org1] and GSM2180145 [Org2]);

H9 embryonic stem cells (Luo et al., Cell Reports 2016): 2 biological replicates (GEO accession numbers: GSM2180138 [ESC1] and GSM2180139 [ESC2]);

Fetal human brain samples (Ataman et al., Nature 2016): 5 biological replicates, unstimulated or stimulated ("stim") with KCl for 6 hours (GEO accession numbers: GSM2072621, GSM2072624, GSM2072627, GSM2072630, GSM2072633, GSM2072623, GSM2072626, GSM2072629, GSM2072632, and GSM2072635).

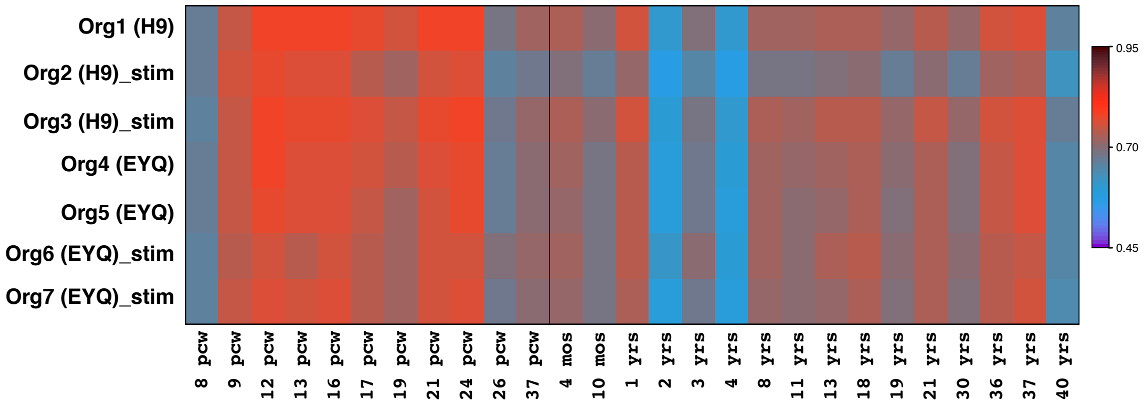

Org1 (H9)
Org2 (H9)_stim
Org3 (H9)_stim
Org4 (EYQ)
Org5 (EYQ)
Org6 (EYQ)_stim
Org7 (EYQ)_stim
8 pcw
9 pcw
12 pcw
13 pcw
16 pcw
17 pcw
19 pcw
21 pcw
24 pcw
26 pcw
37 pcw
4 mos
10 mos
1 yrs
2 yrs
3 yrs
4 yrs
8 yrs
11 yrs
13 yrs
18 yrs
19 yrs
21 yrs
30 yrs
36 yrs
37 yrs
40 yrs
0.95
0.70
0.45

**Supplementary Figure 12. Transcriptional analysis to assess the developmental age of 5-month-old SNR-derived organoids.** Spearman correlation of expression levels between 7 individual 5-month-old SNR-derived organoids and age-specific neocortical human samples obtained from BrainSpan database (http://www.brainspan.org/).

The following neocortical samples from BrainSpan samples were used for analysis: VFC (38 samples at 27 ages), MFC (37 samples at 28 ages), DFC (39 samples at 29 ages). Horizontal axis displays age on a quasi-log scale, beginning at zero post-conception weeks (pcw), through birth (at 40 pcw), up to 50 years; range of ages plotted, 8 pcw to 40 years old.

**a**

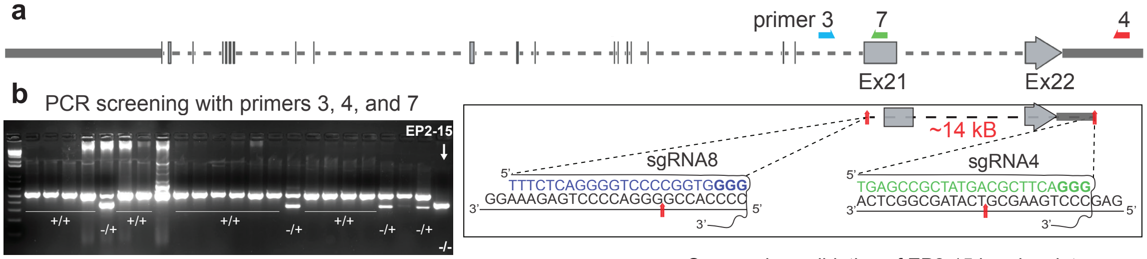


**c** Summary

| Electroporation | Stem cell lines | # colonies selected | # with hetero deletion (+/-) | # with homo deletion (-/-) |
|---|---|---|---|---|
| EP#2 | iPS 2242-5 (P13) | 24 | 4 | **1 (EP2-15)** |
| EP#5 | iPS 2242-5 (P14) | 24 | 0 | 0 |
| EP#16 | ES H9 (P42) | 24 | 4 | 0 |
| EP#19 | ES H9 (P43) | 10 | 1 | 0 |

**d** Sequencing validation of EP2-15 break points

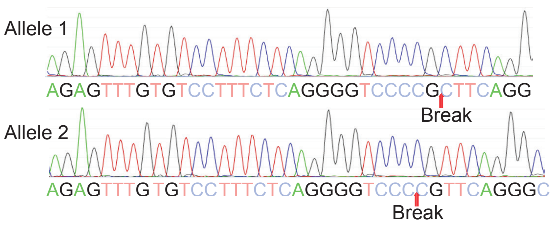


**e** Concensus sequences

2242-5 (wt)

sgRNA8 sgRNA4

AGAGTTTGTGTCCTTTCTCAGGGGTCCCCGGTGGGG..14041bp...TGAGCCGCTATGACGCTTCAGGGCTCTGT

EP2-15 *(SHANK3 -/-)*

AGAGTTTGTGTCCTTTCTCAGGGGTCCCCG--------------(-14062bp)--------------CTTCAGGGCTCTGT

AGAGTTTGTGTCCTTTCTCAGGGGTCCC--------------(-14063 bp)--------------CGCTTCAGGGCTCTGT

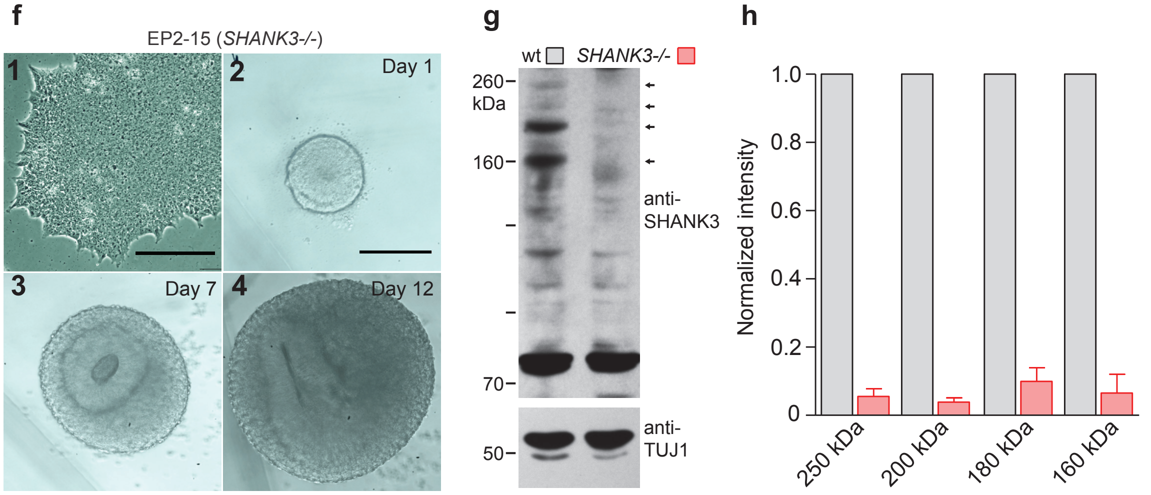

**Supplementary Figure 13. Generation and characterization of CRISPR/Cas9-engineered *SHANK3-/-* iPSC line.**

**a**, Strategy used for introducing *SHANK3* deletion into human iPSCs. Two single-guide RNAs (sgRNA), sgRNA8 and 4, were designed to flank exons 21-22 and to introduce a ~14-kB deletion (zoom in). **b**, PCR verification of the deletion using a set of primers that recognize different regions in the proximity of the exons 21-22. **c**, Homozygous deletion of *SHANK3* was detected in 1 out of 82 tested colonies (EP2-15 clone). **d**, Verification of homozygous *SHANK3* deletion in EP2-15 line using subcloning and sequencing. **e**, Consensus sequences of exons 21-22 region in 2242 (isogenic control) and EP2-15 (*SHANK3-/-*). **f**, Images of iPSCs (**f1**), iPSC-derived neural rosette (**f2**), and SRCOs at different time points after single rosette isolation (**f3-4**). **g**, Images of Western blots of lysates obtained from wild-type (WT) and *SHANK3-/-* neurons immunoblotted using anti-SHANK3 and anti-TUJ1 antibodies. **h**, Quantification of expression of different isoforms of SHANK3 in wt and *SHANK3-/-* neurons (n = 3 pairs of samples). Data are presented as means ± s.e.m. Scale bars = 250 μm.

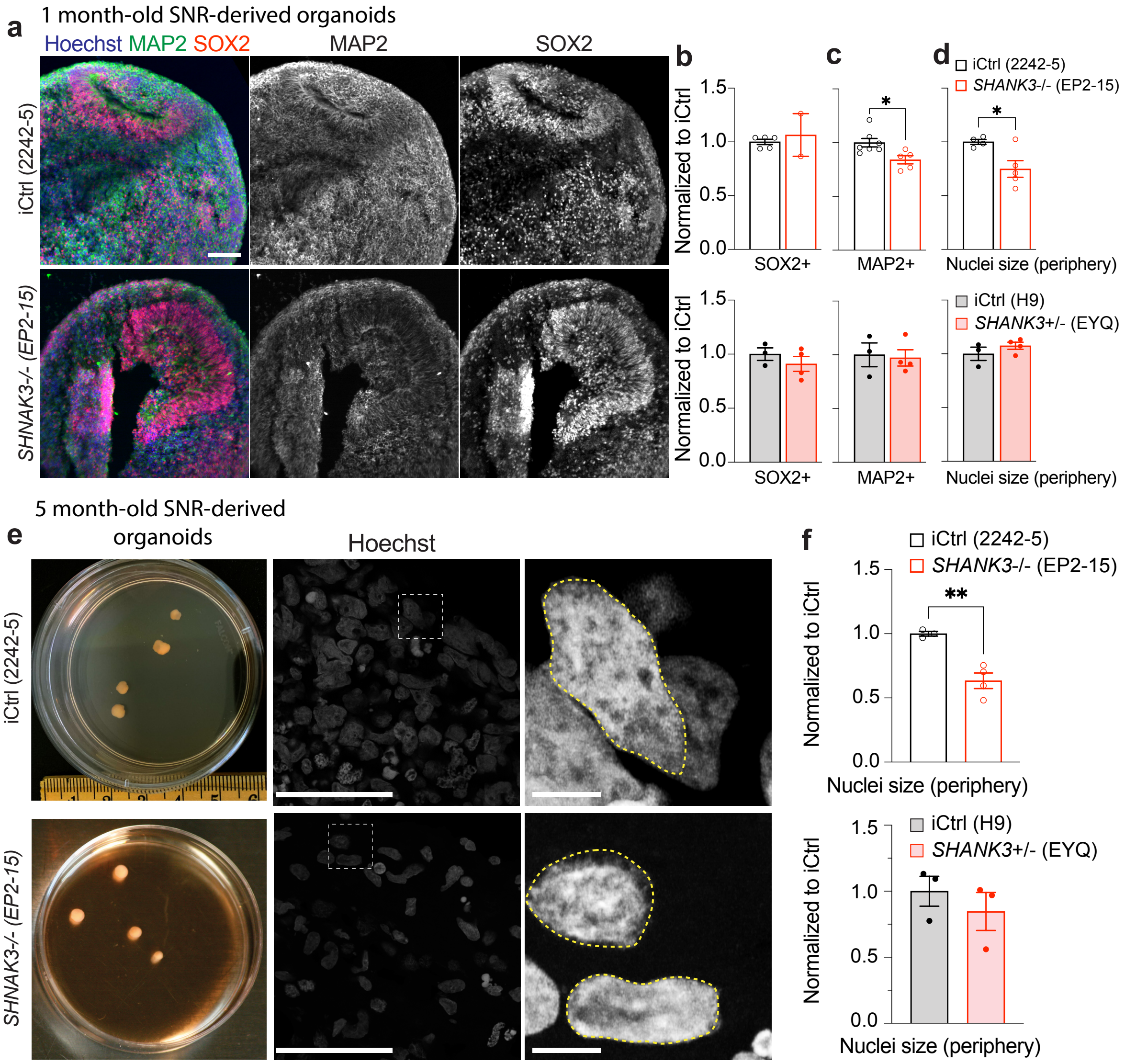

a
1 month-old SNR-derived organoids
Hoechst MAP2 SOX2
MAP2
SOX2
iCtrl (2242-5)
SHNAK3-/- (EP2-15)
b
c
d
iCtrl (2242-5)
SHANK3-/- (EP2-15)
Normalized to iCtrl
SOX2+
MAP2+
Nuclei size (periphery)
iCtrl (H9)
SHANK3+/- (EYQ)
5 month-old SNR-derived organoids
e
Hoechst
f
Nuclei size (periphery)

**Supplementary Figure 14. Characterization of cellular deficits in SHANK-deficient organoids**

**a**, Images of 1-month-old organoid sections from iCtrl (2224-5) and *SHANK3*-/- (EP2-15) organoids stained with anti-SOX2 and MAP2 antibodies and Hoechst. **b-c**, Quantifications of SOX2- (**b**) and MAP2-expressing cells (**c**) in *SHANK3*-/- (top) and *SHANK3*+/- (bottom) organoids as compared to respective iCtrl organoids (SOX2, n = 5 iCtrl [2224-5], 2 *SHANK3*-/- [EP2-15], 3 iCtrl [H9], and 4 *SHANK3*+/- [EYQ2-20] organoids; MAP2, n = 7 iCtrl [2224-5], 5 *SHANK3*-/- [EP2-15], 3 iCtrl [H9], and 4 *SHANK3*+/- [EYQ2-20] organoids). **d**, Quantification of nuclei sizes of neurons near the periphery in *SHANK3*-/- (top) and *SHANK3*+/- (bottom) organoid sections as compared to respective iCtrl organoid sections (n = 4 iCtrl [2224-5], 5 *SHANK3*-/- [EP2-15], 3 iCtrl [H9], and 4 *SHANK3*+/- [EYQ2-20] organoids). **e**, Images of 5-month-old iCtrl (2224-5) and *SHANK3*-/- (EP2-15) organoids and organoid sections stained with anti-Synapsin1 and SHANK3 antibodies and Hoechst. **f**, Quantification of nuclei sizes of neurons near the periphery in 5-month-old SNR-derived organoids (n = 3 iCtrl [2224-5], 4 *SHANK3*-/- [EP2-15], 3 iCtrl [H9], and 3 *SHANK3*+/- [EYQ2-20] organoids). Data presented as mean ± s.e.m.; *P<0.05, **P<0.01, unpaired t-test. Scale bars = 50 (**a, e**) and 5 μm (**e**).

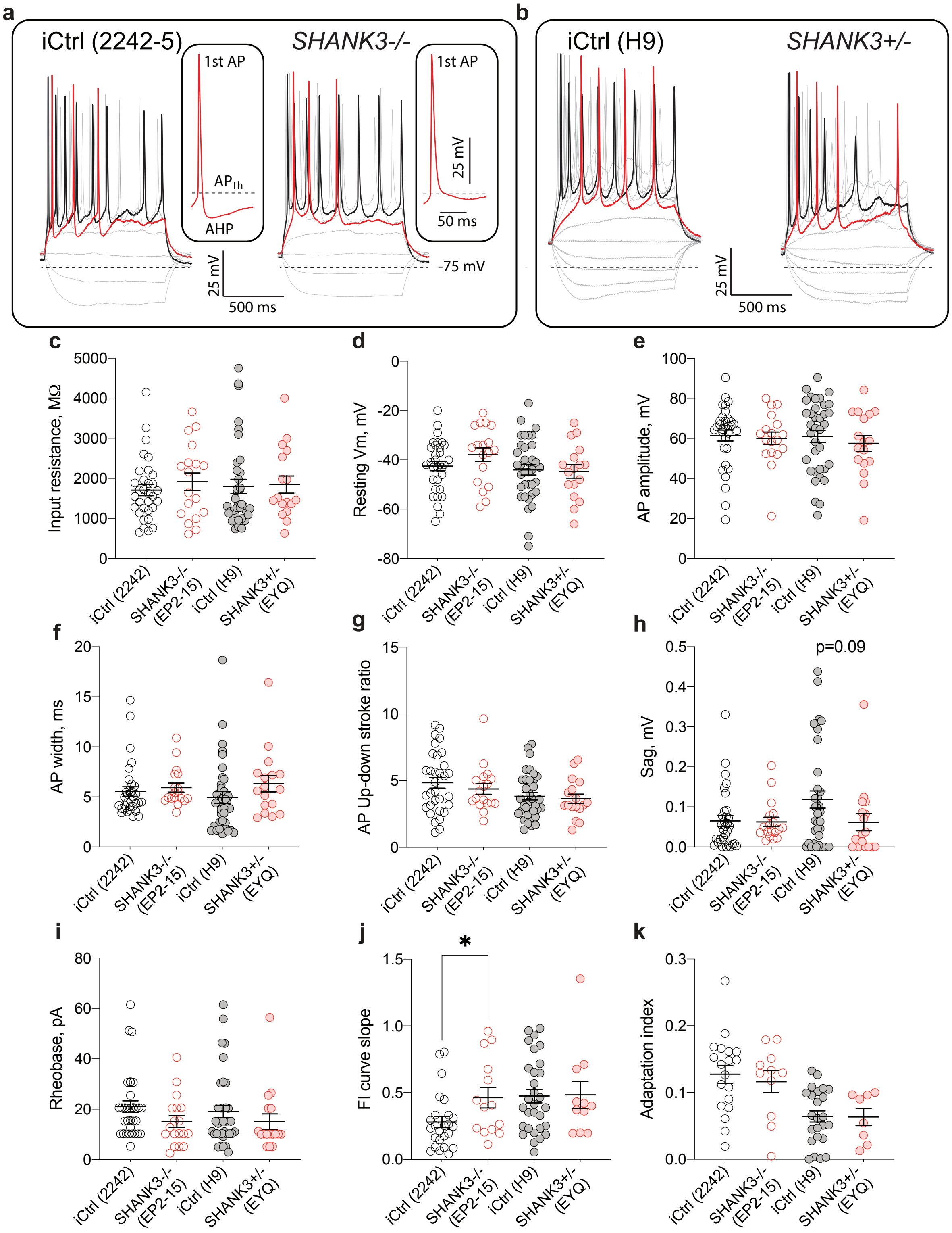
a
iCtrl (2242-5)
SHANK3-/-
1st AP
APTh
AHP
25 mV
50 ms
-75 mV
25 mV
500 ms
b
iCtrl (H9)
SHANK3+/-
c
Input resistance, MΩ
d
Resting Vm, mV
e
AP amplitude, mV
f
AP width, ms
g
AP Up-down stroke ratio
h
Sag, mV
p=0.09
i
Rheobase, pA
j
FI curve slope
*
k
Adaptation index
iCtrl (2242)
SHANK3-/- (EP2-15)
iCtrl (H9)
SHANK3+/- (EYQ)

**Supplementary Figure 15. Characterization of intrinsic properties of neurons in iCtrl and SHANK3-deficient organoids. a-b**, Traces of membrane potentials obtained from SHANK3-/- (**a**) and SHANK3+/- (**b**) neurons as compared to respective isogenic control (iCtrl) neurons in response to different somatic current injections. **c-k**, Quantification of input resistance (**c**), resting membrane potential (**d**), amplitude of the first AP (**e**), width of the first AP (**f**), ratio of the first AP up to down stroke (**g**), sag amplitude (**h**), rheobase (**i**), slope of frequency-current curve (**j**), and adaptation index (**k**). Data presented as individual data points and mean ± s.e.m.; $^*P<0.05$, unpaired t-test.

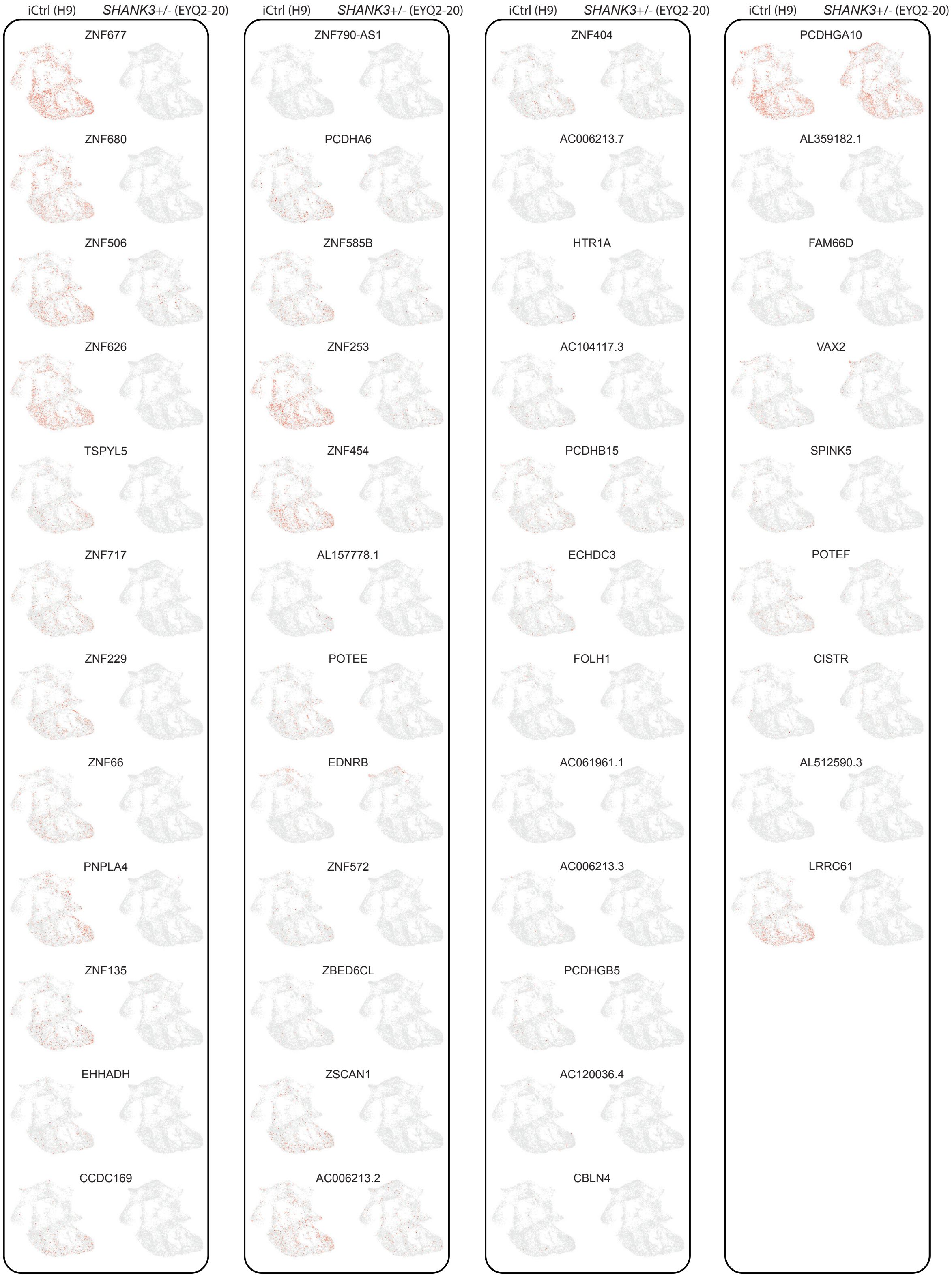
iCtrl (H9)
*SHANK3*+/- (EYQ2-20)
ZNF677
ZNF680
ZNF506
ZNF626
TSPYL5
ZNF717
ZNF229
ZNF66
PNPLA4
ZNF135
EHHADH
CCDC169
iCtrl (H9)
*SHANK3*+/- (EYQ2-20)
ZNF790-AS1
PCDHA6
ZNF585B
ZNF253
ZNF454
AL157778.1
POTEE
EDNRB
ZNF572
ZBED6CL
ZSCAN1
AC006213.2
iCtrl (H9)
*SHANK3*+/- (EYQ2-20)
ZNF404
AC006213.7
HTR1A
AC104117.3
PCDHB15
ECHDC3
FOLH1
AC061961.1
AC006213.3
PCDHGB5
AC120036.4
CBLN4
iCtrl (H9)
*SHANK3*+/- (EYQ2-20)
PCDHGA10
AL359182.1
FAM66D
VAX2
SPINK5
POTEF
CISTR
AL512590.3
LRRC61

**Supplementary Figure 16. scRNA-seq UMAP heatmap plots for 45 most down-regulated DEGs identified using bulk RNA sequencing.** Left columns: iCtrl (H9) organoids (n = 2 organoids). Right columns: Left columns: *SHANK3*+/- (EYQ) organoids (n = 2 organoids).
