## Supplementary Tables for "Modeling autism-associated SHANK3 deficiency using human cortico-striatal organoids generated from single neural rosettes": Suppl Table 7 Primary antibodies.docx

**Supplementary Table: Primary antibodies**

| Antibody target | Company | Catalog # | Raised in | Dilution |
| --- | --- | --- | --- | --- |
| Sox2 | Abcam | ab97959 | Rb | 1:500 |
| Pax6 | BioLegend/Covance | 901301 (previously PRB-278P) | Rb | 1:250 |
| N-Cadherin | Abcam | ab98952 | Ms | 1:500 |
| Ki67 | BD Biosciences | 550609 | Ms | 1:100 |
| Tuj1 | Covance | MMS-435P | Ms | 1:1000 |
| Map2 | Synaptic Systems | 188 004 | Gp | 1:1000 |
| Tbr2 | EMD Millipore | AB2283 | Rb | 1:300 |
| Reelin | MBL International | D223-3 | Ms | 1:500 |
| GABA | Sigma | A2052 | Rb | 1:500 |
| GAD67 | EMD Millipore | MAB5406 | Ms | 1:500 |
| Somatostatin | Chemicon | AB5494 | Rt | 1:500 |
| Parvalbumin | Swant | PV 235 | Ms | 1:500 |
| VIP | Immunostar | 20077 | Rb | 1:500 |
| Calretinin | Swant | 7697 | Rb | 1:500 |
| Calbindin (d28k) | Swant | 300 | Ms | 1:500 |
| S100b | Agilent Technologies | Z031129-2 | Rb | 1:1000 |
| GFAP | Abcam | ab4674 | Ch | 1:300 |
| MBP | Chemicon | MAB386 | Rt | 1:500 |
| O4 | R&D Systems | MAB1326-SP | Ms | 1:500 |
| Tbr1 | Abcam | ab31940 | Rb | 1:500 |
| Ctip2 | Abcam | ab18465 | Rt | 1:500 |
| Satb2 | Abcam | ab51502 | Ms | 1:500 |
| Cux1 | Santa Cruz Biotechnology | sc-13024 | Rb | 1:500 |
| GFP | Abcam | ab13970 | Ch | 1:1000 |
| Caspase-3 | BD Pharmingen | 559565 | Rb | 1:500 |
| PH3 | EMD Millipore | 06-570 | Rb | 1:500 |
| Foxp2 | Abcam | ab16046 | Rb | 1:500 |
| Bassoon | Enzo Life Sciences | ADI-VAM-PS003-D | Ms | 1:500 |
| Homer1 | Synaptic Systems | 160 004 | Gp | 1:500 |
| Gephyrin | Synaptic Systems | 147 003 | Rb | 1:50 |
| Vglut1 | EMD Millipore | AB5905 | Gp | 1:1000 |
| Synapsin1 | Synaptic Systems | 106 001 | Rb | 1:500 |
| PSD-95 | Abcam | ab2723 | Ms | 1:100 |
| Shank1 | Novus Biologicals | NB300-167 | Rb | 1:100 |
| Shank2 | Synaptic Systems | 162 202 | Rb | 1:200 |
| Shank3 | Synaptic Systems | 162 304 | Gp | 1:200 |
| Shank3 | Synaptic Systems | 162 302 | Rb | 1:100 |
| PDGFR-β | Santa Cruz Biotechnology | sc-374573 | Ms | 1:100 |
| FoxG1 (Bf1) | Takara Bio | M227 | Ms | 1:250 |
| Fam107A | Proteintech | 12176-1-AP | Rb | 1:100 |
| αSMA | Thermo Fisher Scientific | 710487 | Rb | 1:100 |

Abbreviations: Rb –rabbit, Ms – mouse, Gp – guinea pig, Rt – rat, Ch –
